# Mutation of sequences flanking and separating transcription factor binding sites in a *Drosophila* enhancer significantly alter its output

**DOI:** 10.1101/379974

**Authors:** Xiao-Yong Li, Michael B. Eisen

## Abstract

Here we explore how mutating different sequences in an enhancer that regulates patterned gene expression in *Drosophila melanogaster* embryos can affect its output. We used quantitative imaging to analyze the effects of a wide variety of mutations in the *hunchback* distal anterior enhancer. This enhancer has been shown to respond to the anterior morphogen Bicoid, but we found that mutations in only one of the five strong Bicoid sites in the enhancer has a significant effect on its binding. The pioneer factor Zelda, which binds to this enhancer and is the only other factor implicated in its activity besides Bicoid. However, we found that mutations of all its sites only has modest effect that is limited to reduction of its output in more posterior regions of the embryo, where Bicoid levels are low. In contrast to the modest effects of mutating known transcription factor binding sites, randomizing the sequences between Zelda and Bicoid sites significantly compromised enhancer activity. Finer mapping suggested that the sequences that determine activity are broadly distributed in the enhancer. Mutations in short sequences flanking Bicoid binding sites have stronger effects than mutations to Bicoid sites themselves, highlighting the complex and counterintuitive nature of the relationship between enhancer sequence and activity.

## INTRODUCTION

Precise spatiotemporal gene expression in animal development is mediated by the binding of sequence-specific, DNA binding transcription factors to cis-regulatory sequences known as enhancers (Lelli et al., 2012; Levine, 2010; Spitz and Furlong, 2012). Since their discovery more than three decades ago (J. Banerji, S. Rusconi, W. Schaffner, 1981) tremendous progress has been made in our understanding of the structure and function of enhancers. However, they remain highly enigmatic: it is difficult to identify enhancer sequences *de novo* based on their sequence alone, let alone predict their transcriptional output.

One of the major, outstanding challenges in understanding enhancers is our limited knowledge of what makes a sequence function as an enhancer. Because transcription factors recognize short, redundant DNA sequences, potential transcription factor binding sites are found throughout animal genomes. Yet only a small fraction of sites for any given factor are bound *in vivo* and are involved in transcriptional regulation(Iyer et al., 2001; Li et al., 2008; Liu et al., 2006; Wunderlich and Mirny, 2009; Yang et al., 2006).

Many hypotheses have been offered to explain why only a small fraction of the sequences containing transcription factor binding sites function as enhancers, including the number of binding sites (Mirny, 2010), the organization and orientation of binding sites to facilitate cooperative interactions between factors (Jolma et al., 2015; Mann et al., 2009; Panne et al., 2007), and various forms of indirect cooperativity (Miller and Widom, 2003; Mirny, 2010). It is now clear that some factors, known as pioneer factors, can potentiate the binding of others (Zaret and Carroll, 2011; Zaret and Mango, 2016) and various, recent studies have also suggested that sequence features other than transcription factor binding sites may play a role in enhancer activity (Slattery et al., 2014).

Here we explore these questions using one of the simpler enhancers active in early *Drosophila melanogaster* development, the distal anterior enhancer (DAE) of *hunchback (hb)* (Perry et al., 2011). The DAE is expressed in a broad anterior pattern in cellular blastoderm (pre-gastrulation) stage embryos, and is believed to be regulated by the anterior morphogen Bicoid (Bcd) (Driever and Nüsslein-Volhard, 1988), which has thirteen sites in the enhancer, and by the maternal pioneer factor Zelda (Zld) (Harrison and Eisen, 2015), which has five. Here we explore the role of these binding sites and other sequences on DAE activity through quantitative imaging of the transcriptional output of several mutations in the DAE.

## RESULTS

To test the effect of mutations on the DAE enhancer activity, we developed a reporter system in which we embedded the enhancer in a bacterial artificial chromosome (BAC) containing the *even-skipped (eve)* gene and all of its regulatory sequences (Venken et al., 2009), as well as the two insulators flanking the *eve* gene locus, which are important for proper function of the enhancers in this locus (Fujioka et al., 2016) (Figure 1A). Our goal was to assay enhancer activity away from its endogenous locus, where other elements might alter its activity, but within the context of a complete gene, rather than using a conventional system with only an enhancer, promoter and reporter gene.

**Fig. 1.**
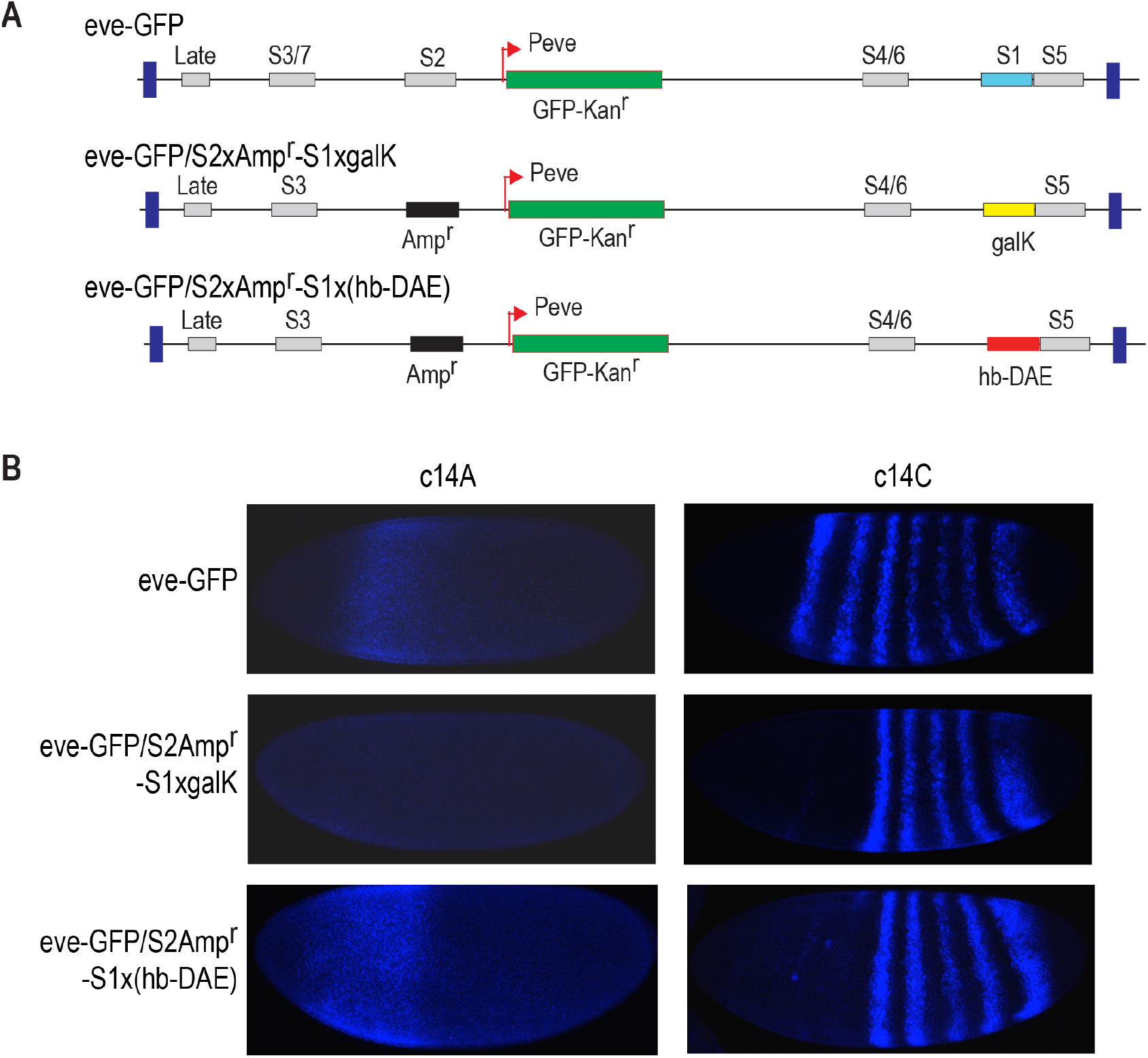
*eve* bac based reporter construct. A). The *eve* bac plasmid contains the full *eve* gene as well as all its regulatory sequences bracketed by two native insulators as shown in dark blue. In the eve-eGFP construct, the CDS of *eve* gene was replaced by the reporter gene eGFP. To create the reporter vector construct eve-eGFP/S2xAmp^r^-S1xgalK, the *eve* stripe 2 and 1 enhancers were further replaced by the bacteria Amp^r^ gene, and the bacteria galK gene, respectively. To create an enhancer reporter construct, e.g. eve-eGFP/S2xAmp^r^-S1x(hb-DAE), the galK sequence in the *eve* bac reporter vector was replaced with the enhancer sequences of interest through recombineering. B) Images showing the reporter activities in early (c14A) and late (c14C) stage 5 embryos from transgenic flies created with the reporter constructs, as detected by FISH using the GFP anti-sense probe.

We first replaced the *eve* coding sequence in the BAC with an eGFP reporter gene, and then replaced the two anterior *eve* stripes, stripe 1 and stripe 2, with galK and Amp^r^. These latter two modifications eliminated any activity from the BAC that overlaps the region where the DAE is normally expressed (Figure 1B). We then replaced galK with the wild-type DAE and various modified versions by BAC recombineering, and generated fly lines containing these constructs. As expected, the construct containing the wild-type DAE expresses in the anterior during early mitotic cycle 14 (c14A) (Perry et al., 2011), and is generally absent after mid-cycle 14 (c14C) (Figure 1B). The activity of the enhancer constructs we report in the remainder of study were all measured in cycle 14A embryos.

### Mutations of Bcd binding sites on *hb* DAE activity

To begin to understand the requirements for the DAE activity, we first focused on the role of potential transcription factor binding sites.

The DAE drives a similar expression pattern to the promoter proximal enhancer of hb(Perry et al., 2011), for which Bcd is known as the primary activator (Tautz, 1988)(Driever and Nüsslein-Volhard, 1989; Struhl et al., 1989). So we expect Bcd is the major activator of the DAE as well. In addition, we found Zld, the maternal factor known to be broadly required for zygotic enhancer function(Harrison and Eisen, 2015; Liang et al., 2008), bound strongly to this enhancer based on previous ChIP-seq data (Harrison et al., 2011), and thus may be important as well. Besides these two factors, we found no other activator in the early embryo transcription network that was consistent with the broad anterior expression pattern displayed by the DAE, with the exception of *hb* itself. *hb* can function as an activator, as was first shown for the *eve* stripe 2 enhancer (Small et al., 1991). However, in a previous ChIP-chip study, *hb* did not bind strongly to the DAE enhancer (Li et al., 2008) and is thus unlikely to contribute to its activity. Therefore, Bcd and Zld are likely to be the main activators of the DAE.

To investigate the role of Bcd binding sites in *hb* DAE function, we started by identifying the Bcd motifs in the DAE sequence using the patser program(Hertz and Stormo, 1999). A low stringency cutoff in the motif search was used to ensure that all potentially functional Bcd motifs were identified. For comparison, using the same cutoff parameters, we were able to identify all previously identified Bcd sites in two well studied enhancers including the *eve* stripe 2 enhancer (Small et al., 1992) and the *hb* promoter proximal enhancer (Driever and Nüsslein-Volhard, 1989), as well as multiple additional weaker sites: three for *eve* stripe 2, and five for the *hb* proximal enhancer. This demonstrates the cutoff we used is more than adequate to identify all functionally significant Bcd sites.

We made six single and double mutations covering five these motifs, including the four strongest motifs found in this enhancer sequence, and tested their effect on the reporter activity in the embryos of their corresponding transgenic flies (Fig. 2A). We found that among all the mutant enhancers tested, only B1 single site mutated enhancer and the B1 and B2 double site mutated enhancer caused significant decrease in the enhancer activity compared to the wild type enhancer. The activities of these two mutant enhancers are very similar, which, combined with the observation that the other two double site mutants involving B2 did not have significant effect (Fig. 2B), suggests that mutation of the B1 site is mainly responsible for the effect observed for the B1 and B2 double mutant, and the B2 site does not seem to play a important role in DAE activity. Taken together, among the five Bcd sites mutated, only one site, B1, showed significant effect on the DAE activity. This result is very surprising, and suggests that this enhancer harbors an excess of Bcd sites, which presumably is important to ensure robustness of the enhancer against mutational perturbation.

**Fig. 2.**
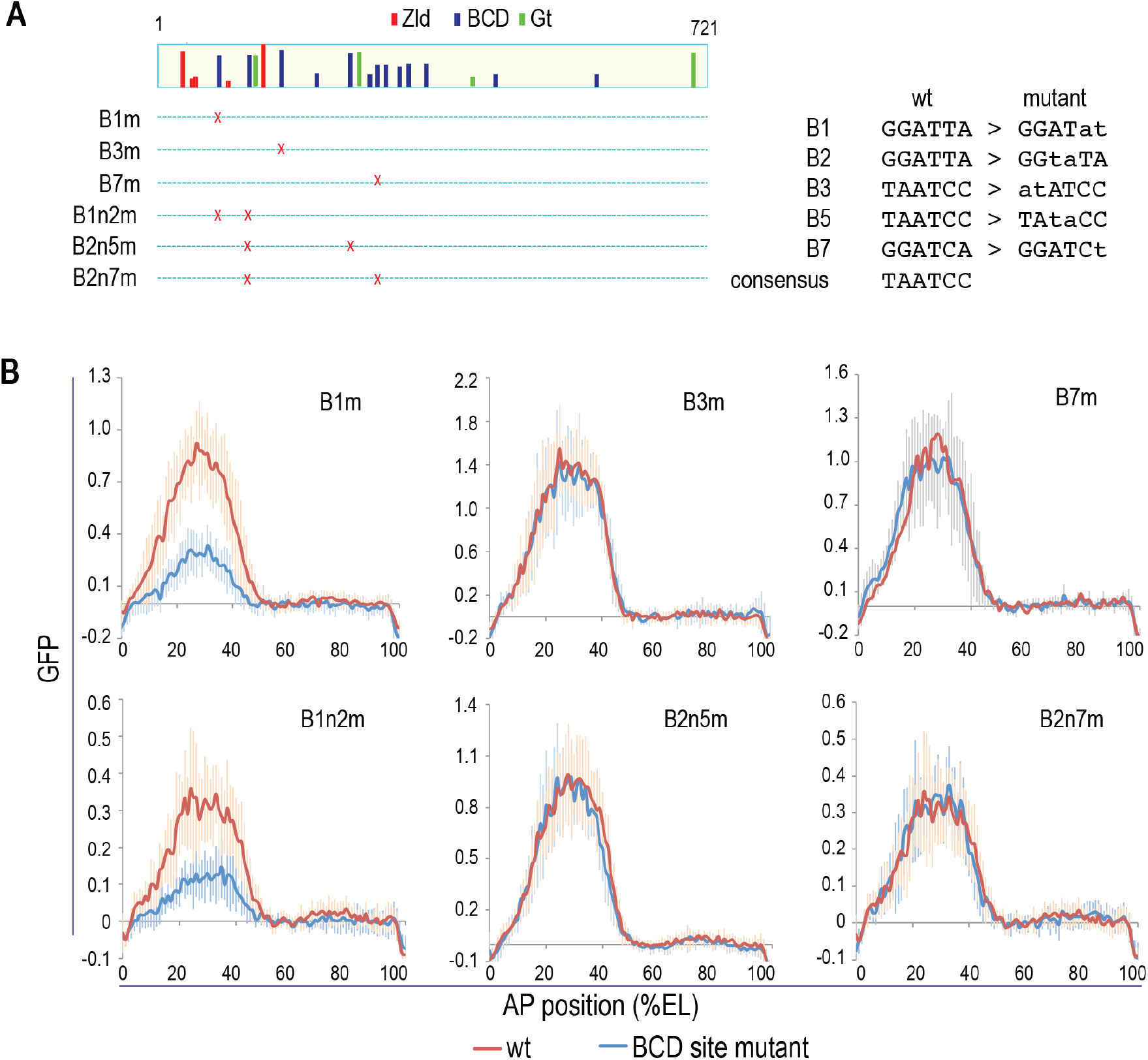
Effect of Bcd binding site mutations. A) The distributions of Bcd, as well as Zid and Gt motifs in the *hb* DAE sequence are shown schematically; the heights of the bars correspond to the motif weight matrix scores. The single or double Bcd binding site mutations are as shown below. The Bcd sites were named based on their order in the enhancer sequence. B) The activities of the mutant enhancers. The GFP reporter activity in embryos of the transgenes was detected by FISH using a GFP antisense probe, and the GFP signal across the A/P axis was quantified and normalized with nuclei density, as described in Material and Methods.

### Zid is only partially required for *hb* DAE activity and fails to convert Bcd binding site enriched sequences into enhancers

We next investigated the effect of Zld binding sites on the DAE activity. Using patser at low cutoff, we identified five Zld motifs in the enhancer sequence. We, then created a mutant enhancer with mutations at all these motifs, except the one that is very weak (Fig. 3A). Surprisingly, we found that these mutations had relatively modest effect on the DAE activity (Fig. 3B), and interestingly, while the mutant enhancer displayed significant lower activity toward the posterior part of the DAE expression pattern, there was no effect in the anterior part of the embryo (Fig, 3C). The lack of general requirement for Zld for the activity of this enhancer is surprising, but consistent with a model in which Bcd enhancers active further away from the embryo’s anterior are often more dependent on Zld due to the presence of low Bcd concentration in those cells in the Bcd morphogen gradient, while those that are active close to the anterior, where Bcd concentration is high, are less dependent (Hannon et al., 2017; Xu et al., 2014).

**Fig. 3.**
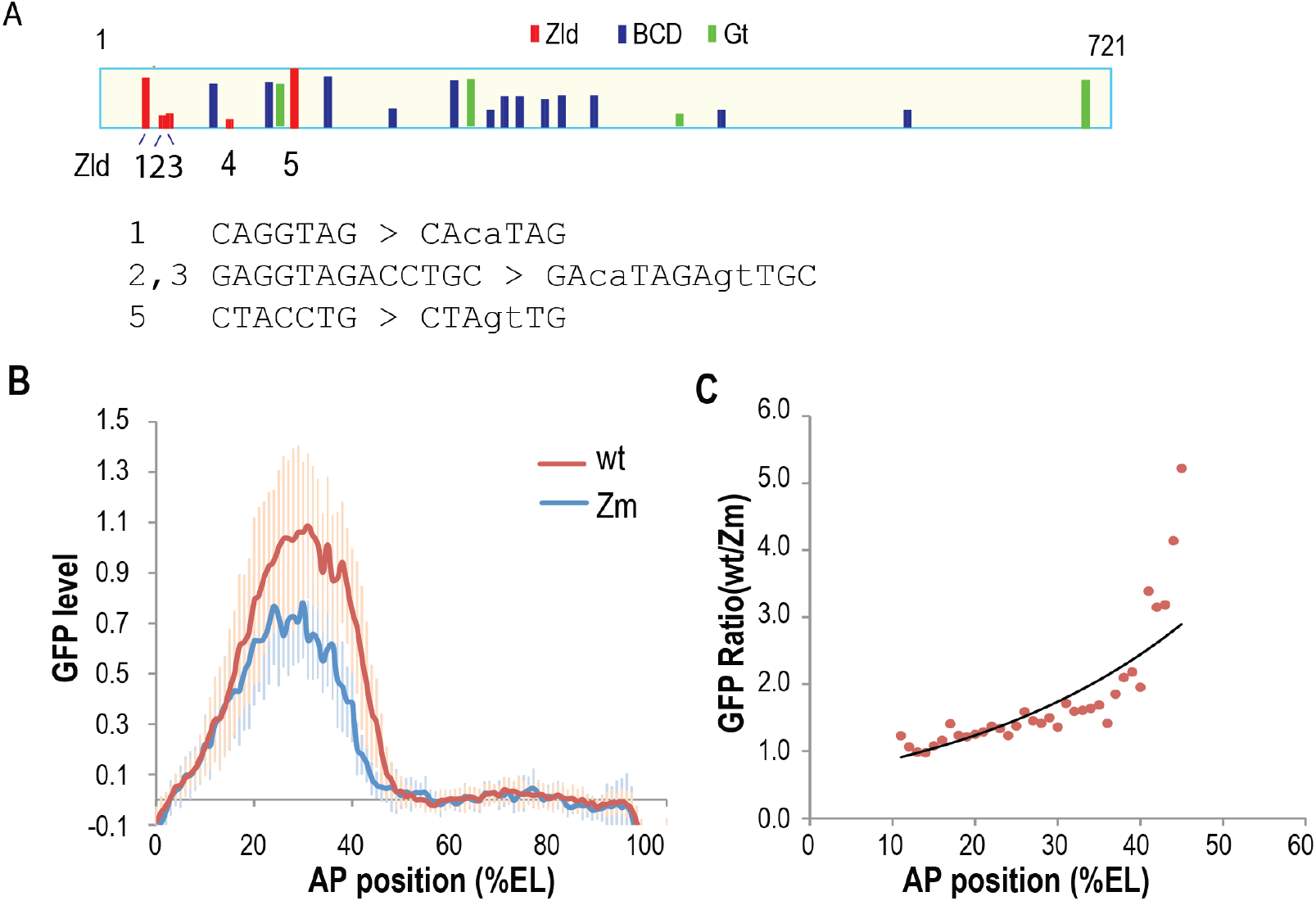
Effect of Zld binding site mutation. A) Five Zld sites are found in *hb* DAE enhancer sequences. All except the weakest site were mutated as shown. B) The reporter activity detected in transgene embryos for the Zld binding sites mutant enhancer is significantly lower than measured for the wild type enhancer. The GFP reporter activity was assayed, quantified and normalized as described Fig.2B. C) The ratio of the reporter activity of the wild type enhancer to activity of the Zld binding site mutant enhancer is plotted against the relative distance from anterior pole of the embryo, with the distance expressed as percentage of egg length (EL).

The seemingly simple requirement for the DAE activity, just a mere combination of Bcd and Zld sites, is very interesting since there are a large number of genomic regions that are enriched with Bcd binding sites but do not function as enhancers and are not bound by Bcd *in vivo* based on ChIP studies (Xu et al., 2014). We wondered whether Zld can convert these sequences enriched for Bcd sites, which presumably contain less than the adequate number and arrangement of Bcd sites, to active enhancers.

To test the above possibility, we selected two genomic sequences that have six and eleven Bcd sites respectively, and cloned them into the reporter vector with or without additional Zld sites added to the sequences. We inserted the Zld motif containing sequences from *hb* sites into the sequences in such a way so as to make the arrangements of Zld motifs relative to Bcd motifs somewhat similar to those present in the *hb* DAE (Fig. S1A). It is worth noting that even though one of these sequences has fewer Bcd sites than *hb* DAE, based on our analysis of known Bcd target enhancers, we found that *hb* DAE possesses the highest number of Bcd sites, and many enhancers have, and can function with, relatively few Bcd sites. After reporter activity was analyzed in the embryos of transgenic flies created with these constructs, we found that neither the original sequences nor the derivatives with the Zld site added displayed any activity (Fig. S1B and S1C). This adds to a list of failed examples from previous attempts to convert genomic sequences enriched with Bcd motifs into active enhancers (Xu et al., 2014), even though a combination of limited number of Zld sites and other activator binding sites was able to activate reporter expression in simpler constructs in which those binding sites were placed direct upstream of a promoter that drives the reporter expression(Crocker et al., 2017).

### The DNA sequences between transcription factor binding sites are crucial for *hb* DAE enhancer activity

The results presented above as a whole are very puzzling: on the one hand the activity of the native enhancer was only modestly affected or not affected at all by mutations in its transcription factor binding sites, while on the other hand, adding strong Zld binding sites to sequences that would appear to contain the adequate number of Bcd sites failed to convert them into active enhancers. While this may be due to lack of proper arrangement of the motifs in the non-enhancer sequences, among other possibilities, we contemplated the possibility that enhancer sequences besides transcription factor binding sites may also be important for enhancer activity.

**Fig. S1.**
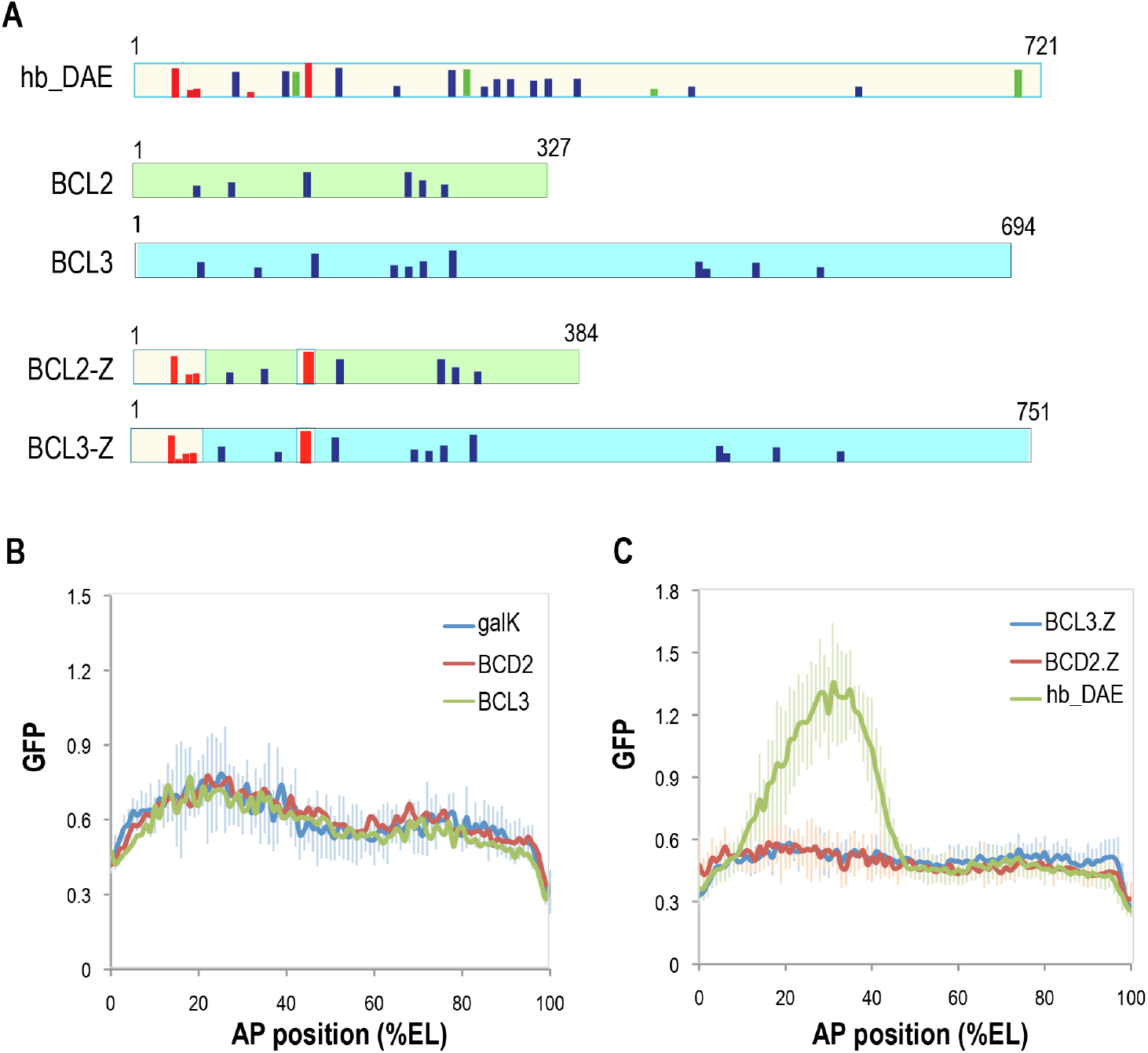
Adding Zld binding sites to Bcd site-enriched non-enhancer sequences failed to convert them into active enhancers. Two of the highly ranked regions, Bcl-2 and Bcl-3, based on Bcd site enrichment, either without or with the addition of Zld binding sites, were tested by cloning into the *eve* bac reporter vector. A. Distribution of the Bcd sites and the added Zld sites are shown. B. The GFP reporter activities in the embryos from the transgenes of the Bcl-2, 3 containing bac reporter constructs. The reporter activity was assayed, quantified and normalized as described Fig.2B. C. Similar to B, except the GFP reporter activities in the embryos from the transgenes of the bac reporter constructs of Bcl-2, 3 sequences with Zld sites added are shown.

To investigate the above hypothesis, we decided to test the effect of mutating sequences around transcription factor binding sites. As mentioned above, Bcd and Zld are likely to be the major, if not sole activators of the DAE. In addition, based on ChIP-chip studies (MacArthur et al., 2009), this enhancer is also bound by Giant (Gt) and Tailless (Tll). These two factors are repressors, and thus are of lesser significance in our analysis. Nevertheless, we took the Gt sites into consideration in our mutational analysis.

We created two mutant enhancers (Fig.4A). For the first, hb-DAE(Rd), we kept the core motif sequences of Zld, Bcd and Gt in the enhancer and replaced the sequences in between with randomly selected sequences of same length. For the other, hb-DAE(TBS), the non-transcription factor binding site sequences were altered by swapping the top and bottom strand bases for each base pair. The mutant enhancers were inserted into *eve* bac reporter and their activities were tested in the embryos of their transgenic flies. Surprisingly, these mutant enhancers displayed dramatically decreased activity compared to the wild type - with up to about thirty fold decrease for hb-DAE(Rd) and about eight fold for hb-DAE(TBS) (Fig.4B-E). This result suggests that the sequences between the transcription factor binding sites are important for *hb* DAE enhancer activity.

**Fig. 4.**
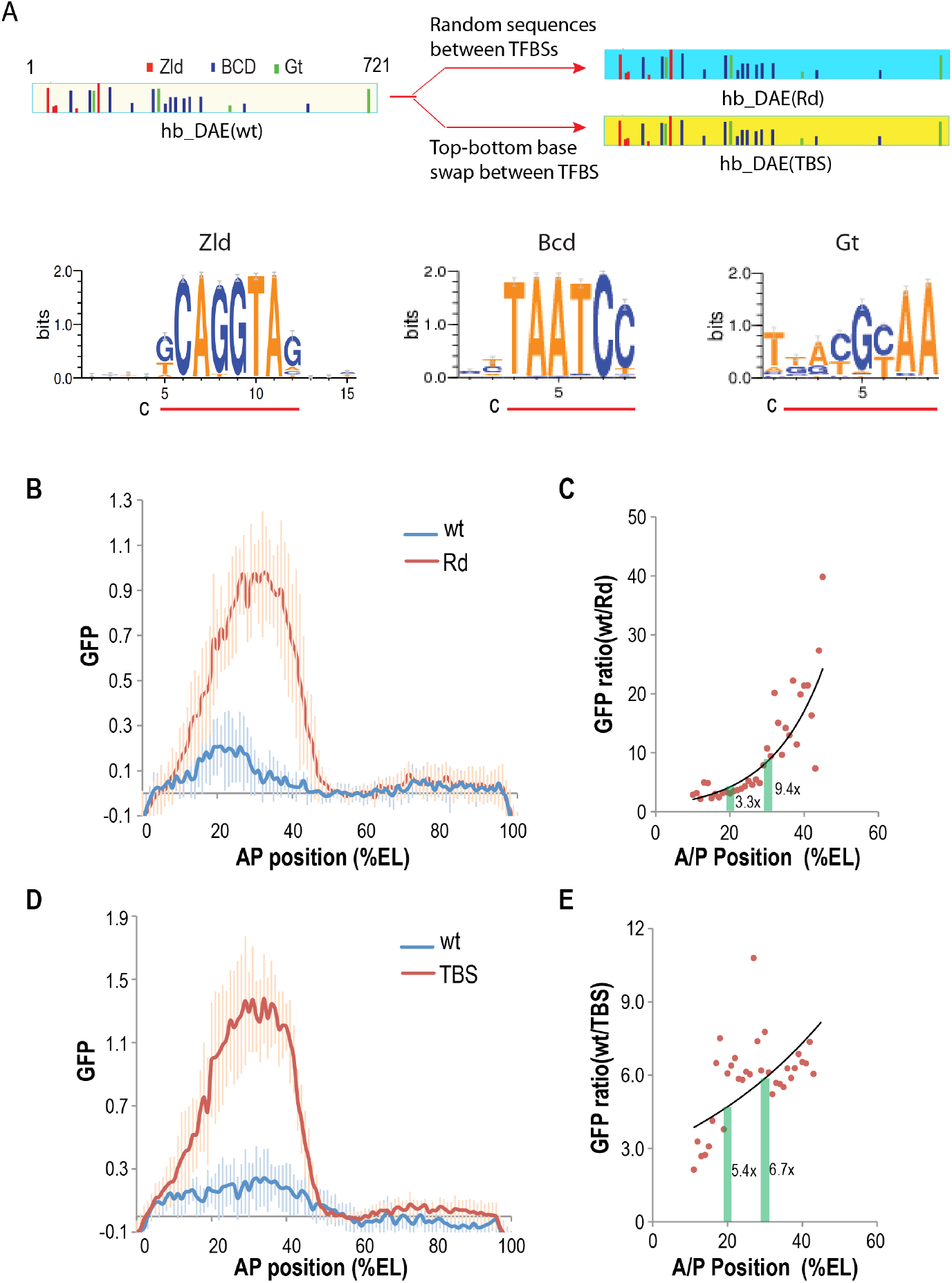
Effect of changes in sequences between transcription factor binding sites on *hb* DAE activity. A). Mutant *hb* DAE enhance sequences were created either by replacing the sequences between transcription actor binding sites that correspond to the core motifs (c) of Zld, Bcd, and Gt with a set of random sequences of lengths equal to the sequences replaced (hb-DAE(Rd)), or by altering the sequences by swapping the top and bottom bases for each base pair (hb-DAE(TBS)). B,D) The GFP activities in the embryos with transgenes of the reporter constructs containing the wild type and mutant *hb* DAE enhancers were assayed, quantified and normalized as described Fig.2B. C,E) The number of fold by which the reporter activity decreased as a result of mutations of the enhancer sequences at increasing distance, by egg length, from the anterior of the embryo.

To see whether the importance of non-transcription factor binding sequences is limited to the *hb* DAE, we carried out similar analysis with the *eve* stripe 2 enhancer (Goto et al., 1989; Small et al., 1992; Stanojevic et al., 1991). We chose to use an expanded version of the enhancer that include additional sequences upstream and downstream of the previously defined minimal stripe 2 enhancer (Small et al., 1992). This enhancer is bound by more transcription factors, which include Bcd, Hb, Kruppel (Kr), Gt, and Zld based on previous studies and a published and ChIP-seq data (Harrison et al., 2011; Small et al., 1992). In addition, previous ChIP-chip data suggests that Tll, the terminal repressor, may also be important for *eve* stripe 2 patterning (Li et al., 2008), but was not considered in the following experiment. Again using the patser program, we identified the binding sites of these factors. For Bcd, Hb, Kr, and Gt the identified motifs included all the sites revealed by previous biochemical and mutational studies(Small et al., 1992), with the exception of two Gt sites that were identified by sequence match to the consensus motif but were shown to have no effect on *eve* stripe 2 enhancer activity when mutated(Arnosti et al., 1995). In addition, other, weaker motifs for Bcd and Kr that were not described in previous studies were also identified. Thus, we are confident that we identified all the binding sites for these factors, and likely for Zld as well, even though no information for Zld on this enhancer was previously available.

We have created mutated *eve* stripe enhancers either by replacing the sequences between the transcription factor binding sites with random sequences (eve-S2(Rd)) or swapping the top and bottom bases for each base pair of the sequences between the sites (eve-S2(TBS)) as we did for the *hb* DAE. Analysis of the reporter activities in the embryos from the transgenes of these mutant enhancer constructs showed that the activity of the *eve* stripe 2 enhancer is completely abolished after these changes (Fig. S2). At the same time, the eve-S2(Rd) displayed a broad domain of expression in the anterior part of the embryo. This ectopic activity may be explained by the fact that the binding sites of Tll, which are likely to be largely responsible for repressing *eve* stripe 2 enhancer activity in the anterior part of the embryo, were mutated in this mutant enhancer. The strong activity in the anterior part of the embryo was not observed for the TBS mutant enhancer. This can be explained by the fact that Tll motif is dyad-symmetric, and as a result the top/bottom strand base conversion often conserved the Tll binding motif. In all, these results showed that the alteration of sequences between the transcription factor binding motifs had a strong, detrimental effect on *eve* stripe 2 enhancer activity. The fact that the sequence randomization mutant displayed enhancer activity in the anterior part of the embryos suggests that the randomization only weakened but did not abolish the capacity of the *eve* stripe 2 enhancer to function. This weaker effect compared to what was observed for *hb* DAE mutants may be due to the presence of large number of transcription factor motifs present in this enhancer and as a result, a smaller percentage of the enhancer sequence was changed. Nevertheless, this result showed that the sequences between transcription factor binding sites are important for *eve* stripe 2 enhancer as well. This result is similar to a previous report, in which a synthetic enhancer containing all transcription factor binding sites for *eve* stripe 2 failed to activate transcription(Vincent et al., 2016).

**Fig. S2.**
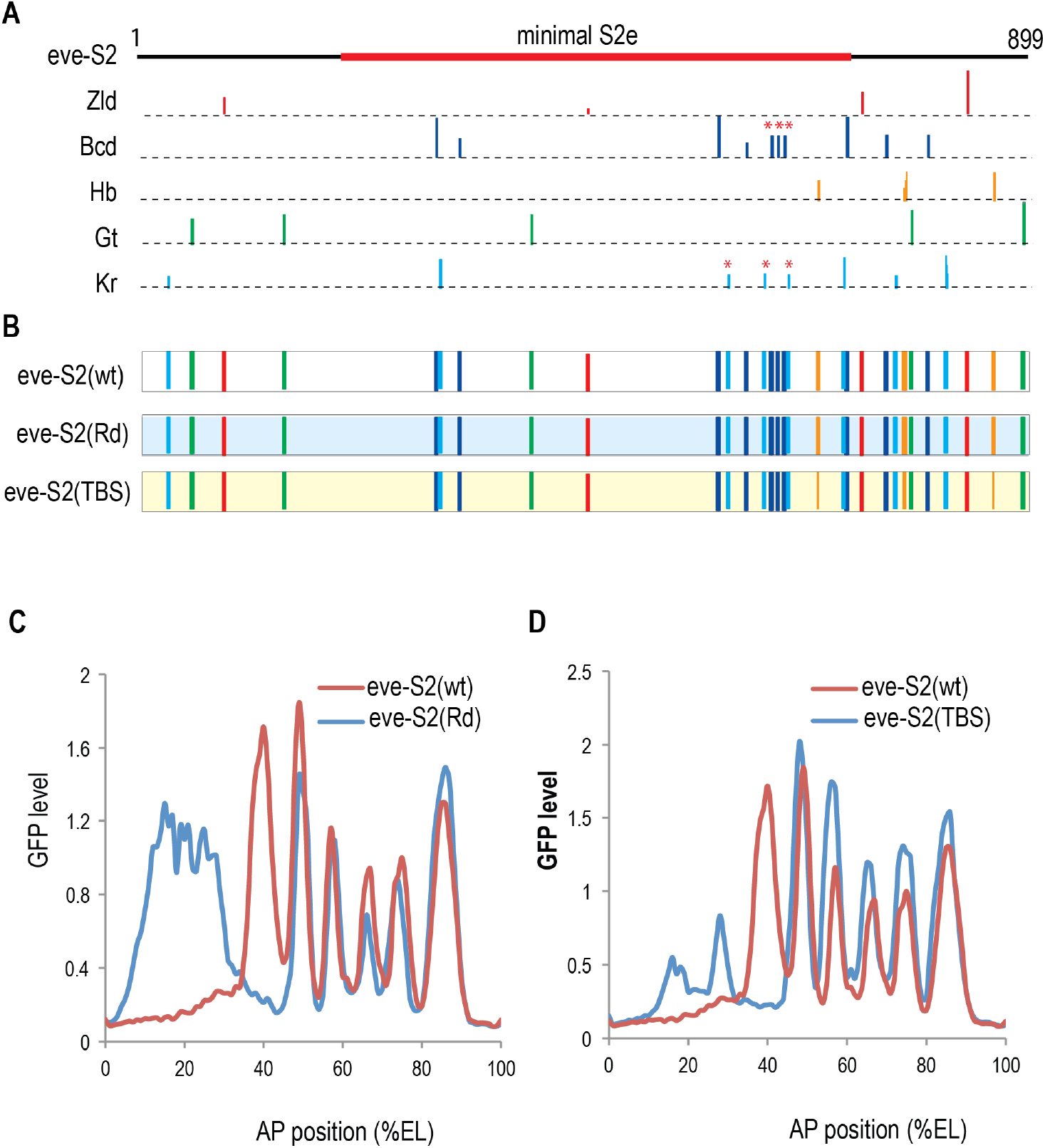
The effect of altering sequences between known transcription factor binding sites on *eve* stripe 2 enhancer activity. A. Schematic representation of the *eve* stripe 2 enhancer. The binding sites for Bcd, Hb, Kr, and Gt identified by patser are as shown, with those identified by patser but not by previous biochemical studies marked by *. The enhancer sequence which we used in our analysis is larger than the minimal *eve* stripe 2 enhancer (Arnosti et al., 1995) defined previously and contains additional factor binding sites as shown. B. To create the mutant enhancers, eve-S2(Rd), and eve-S2(TBS), the sequences between the transcription factor binding sites were altered as described in Fig. 4 for *hb* DAE. C. The GFP reporter activity in mid-stage 5 embryos from the transgenic flies harboring the bac reporter construct, in which the eve-S2(Rd) mutant enhancer was cloned in place of the *eve* stripe 1 enhancer is shown. The reporter activity was assayed, quantified and normalized as described Fig.2B. D. The same as C, except the activity of the reporter construct containing the eve-S2(TBS) mutant enhancer is analyzed.

### The non-transcription factor binding site sequences important for DAE activity are broadly distributed

One possible explanation for the observed effect caused by mutations in the sequences between known transcription factor binding sites is the existence of in those sequences of binding sites for certain known or unknown factor(s). If this is true, it would be possible to identify them by mutating smaller segment of the DAE.

To investigate the above possibility, we divided the 5’ part of the DAE sequence into ~200 bp regions, and replaced the sequences between transcription factor binding sites in each of these regions with random sequences and kept the rest of the enhancer sequence unchanged (Fig. 5A). We found that mutations in first two regions strongly decreased the enhancer activity (Fig. 5B). The sequence changes in the 170 bp region that followed had a relatively small positive effect, leading to an expansion of the expressed domain (Fig.5B), which probably is due to the mutation of certain repressor binding site(s) located in the region. It is worth noting that the first 400 bp of the DAE sequence already contains most of the transcription factor binding sites, so the small effect observed for the mutations in the 170 bp region is not surprising.

**Fig. 5.**
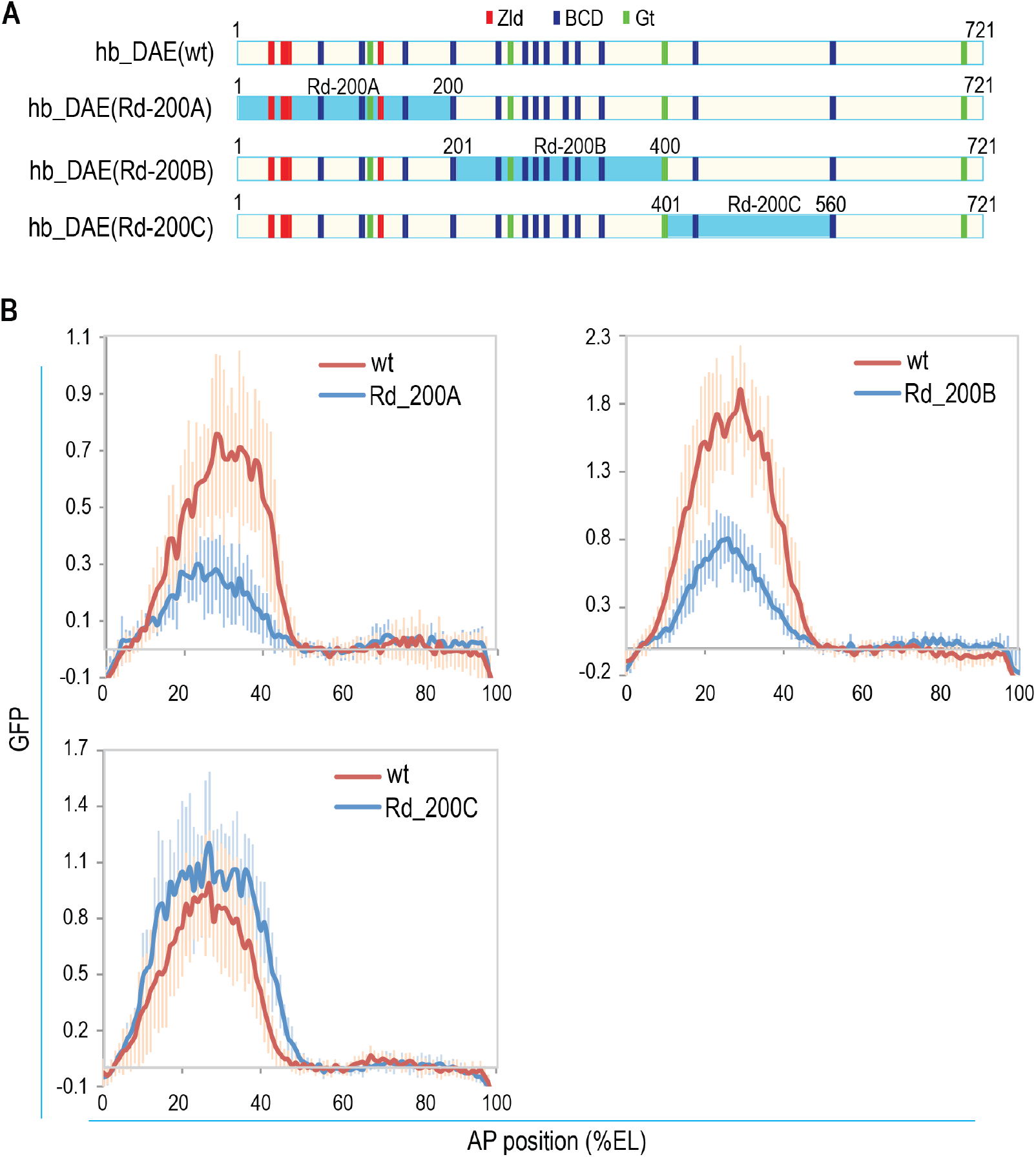
Effect of replacing the sequences between transcription factor binding sites in 200 bp regions in *hb* DAE enhancer on its activity. A) The regions of enhancers in which the sequences between transcription factor binding sites were replaced with random sequences, which corresponds to the sequences in the hb_DAE(Rd) (Fig. 4), are shown. B) The GFP reporter activities of mutants and the wild type *hb* DAE enhancer were assayed, quantified and normalized as described Fig.2B.

To further identify the sequences that may be important, we made four consecutive 50 bp randomization constructs located between positions 77 – 290 (Fig. 6A). Analyses of the reporter activities in the transgenes of these constructs showed that three of the four, Rd-50A-C, had similar strong effect while the fourth, Rd-50D, had a small but significant effect (Fig. 6B). These results again suggest that the non-transcription factor binding site sequences important for *hb* DAE activity are broadly distributed. These results are also surprising when compared to the effects of Bcd binding site mutations described above. In particular, in the three regions 50A-C, there are four Bcd sites, with two in 50A, one in 50B, and one in 50C. As shown above, among these Bcd sites, only mutation of B1 in 50A decreased the DAE activity, while mutations in of B2, also in 50A, and B3, the only Bcd site in region 50B had no effect. The 50C region has only one weak Bcd site, which is probably dispensable for DAE activity as well if mutated. Taken together, these results demonstrate that the sequences between Bcd binding sites are broadly required for DAE activity, and significantly, the changes of these sequences in small regions can have strong effects on the DAE activity even as mutations of Bcd sites in those regions often do not.

**Fig. 6.**
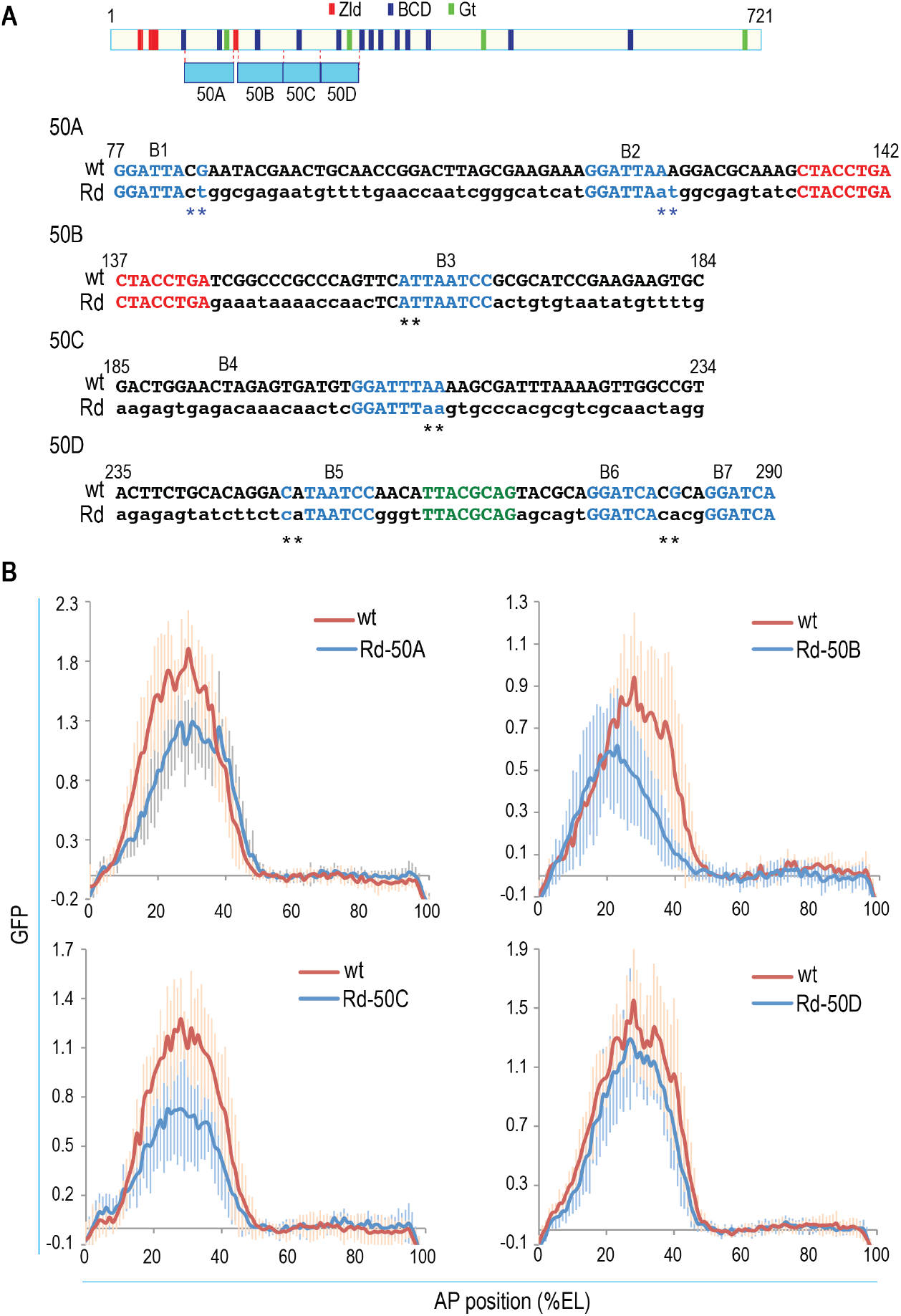
Effect of replacing the sequences between transcription factor binding sites in 50 bp regions in the *hb* DAE enhancer. A) The regions of enhancers in which the sequences between transcription factor binding sites were replaced with the sequences from hb_DAE(Rd) at the corresponding regions, is shown. The wild type sequence and the corresponding mutant sequence in each region are also shown, with the Bcd and Zid binding sites marked in blue and red, respectively. The two base pairs upstream of the BCD core motifs are indicated by star symbols under the sequences. The preferred bases, based on the Bcd binding matrix, are also marked in blue. B) The GFP reporter activity of mutant and the wild type enhancers was detected, quantified and normalized as described in Fig.2B.

### The importance of the non-TFBS sequences can only be partly attributed to the weakly preferred bases flanking the Bcd core motifs

The flanking sequences of the core motifs of transcription factors can have a significant effect in modulating their binding affinity and specificity (Farley et al., 2015; Gordân et al., 2013; Levo et al., 2015). The extent of the effect, as revealed in previous studies, depends on the factor and how the “core” motif is defined. Bcd has two weakly preferred base pairs upstream of the “core motif” in its full DNA binding matrix (Fig. 4A). The flanking sequences of the Zld “core” motif are longer, but the preference is extremely weak (Fig. 4A). Importantly, relatively few bases upstream of the Bcd motifs were changed from the wild type in the hb-DAE(Rd) mutant construct, and among those with changes, the number of sites where favored bases were changed into unfavored bases based on the Bcd binding site is similar to the number of instances where unfavored bases were changed into favored ones (Fig. S3). In addition, in the 50B and 50C mutant constructs which had lower activity than the wild type DAE, the two flanking bases pairs of the Bcd sites are the same as in the wild type (Fig. 6A). Therefore, lower activities of 50B and 50C mutant are not due to changes of the two flanking base pairs of the Bcd motifs in those regions.

**Fig. S3.**
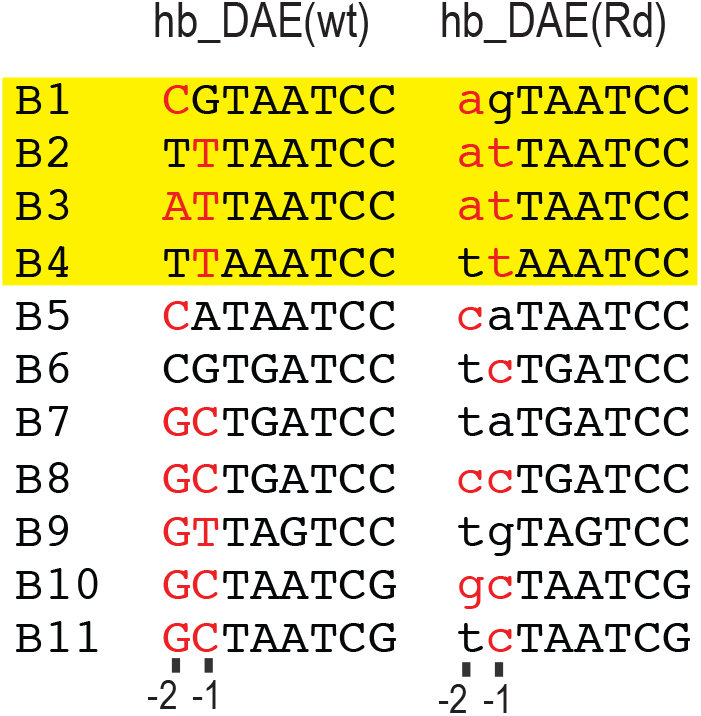
The nucleotide identities at the two proximal positions upstream of the core Bcd binding sites in wild type and mutant *hb* DAE. The preferred bases according to the full Bcd binding matrix are marked in red. The four sites, B1-4, located in the three 50 bp regions, 50A-C, for which the changes in the sequences between the core transcription factor binding sites significantly decreased the enhancer activity are highlighted.

To more critically assess the role of the sequences between the transcription factor binding sites in the enhancer activity and address the contribution of the weakly preferred bases around the Zid and Bcd binding sites, we created a new construct hb-DAE(Rd-f), in which only the sequences between the “full” Bcd, Zid, and Gt motifs were replaced with the corresponding random sequences from hb-DAE(Rd (Fig. 7A. Fig. 7B shows that this mutant enhancer displayed higher activity then the original hb-DAE(Rd) (Fig. 7B). However, its activity is still at least 2 fold less than to the wild type, which provides further evidence for the importance of sequences between transcription factor binding sites on the DAE activity.

**Fig. 7.**
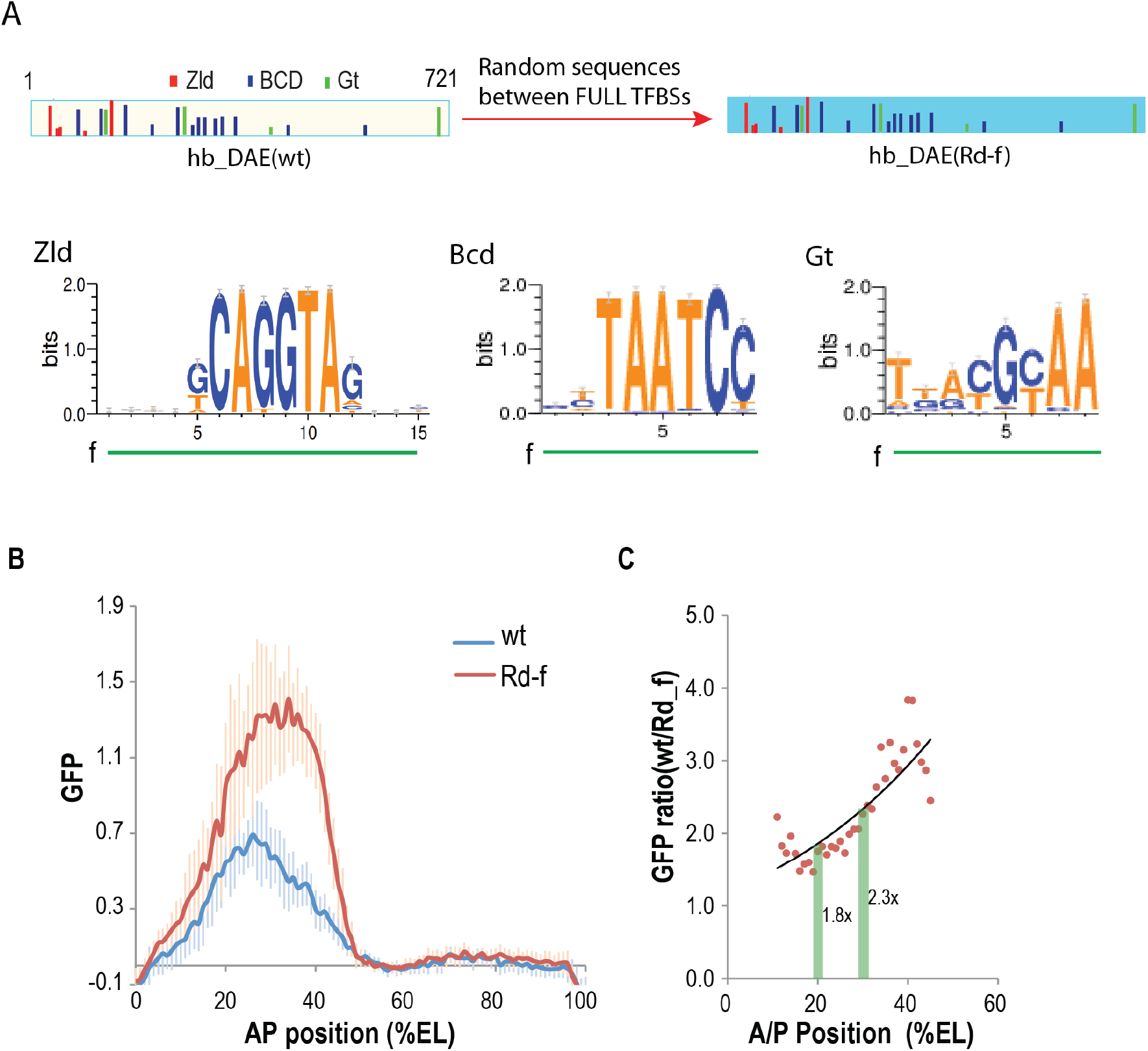
Effect of replacing the sequences between full transcription factor binding sites in *hb* DAE enhancer. A. The mutant enhancer hb-DAE(Rd-f) was made by replacing the sequences between full Zid, Bcd, and Gt binding sites with random sequences, essentially restoring the flanking sequences around the core motifs in the hb-DAE(Rd) to the sequences of the wild type. B. The GFP reporter activity of mutant and the wild type enhancers, assayed, quantified and normalized as described Fig.2B. C). The magnitude of effect caused by the mutations at increasing distance from the anterior pole of the embryo.

### Sequence alignment between different drosophila species reveals conserved sequence blocks

The results shown above demonstrate that sequences between Zld, Bcd, and Gt sites are important for the DAE activity and the effect is broadly distributed. This suggests that the mutational effect probably is not due to the loss of binding site(s) for a certain factor, since if such factor existed, it would need to be able to bind to, and consequently have binding site present in each of the 50 regions, which we find is not true.

Regardless whether factor binding is important between the known transcription factors for the DAE activity, analysis of sequence conservation between different *Drosophila* species can aid the identification of potential transcription factor binding sites and other sequences important for enhancer activity. We therefore carried out sequence alignment for *hb* DAE sequences from the twelve sequenced *Drosophila* species. This analysis revealed conserved blocks (Fig. S4–5). Most these conserved sequences correspond to binding sites for Zid, Bcd, and Gt (which were taken into consideration in the current study and two other known factors, Tll and Knirps (Kni) (Li et al., 2008). Both Tll and Kni are repressors, thus mutation of their binding sites were not expected to negatively affected enhancer activity as we observed by mutating the sequences between Zld, Bcd, and Gt.

There are several other highly conserved, unknown sequence blocks (un1-4) between the transcription factor binding sites. Some extensively conserved sequences were also found around several of the Bcd binding sites. These conserved sequences do not correspond to the preferred dinucleotides upstream of the Bcd core motif and differ between different Bcd motifs. We attempted to find matches for these conserved sequences to known motifs in the combined fly factor database by TomTom (http://meme-suite.org/tools/tomtom) but were unable to find any matches. Besides this lack of match to known motifs, it is also important that none of the conserved sequence motif that appeared more than once and could potentially explain the broad distribution of the mutational effect as discussed above. Thus, different types of sequences between Zld, Bcd, and Gt binding sites may be important for the *hb* DAE activity. It is likely that most if not all of these conserved blocks do not function as transcription factor binding sites.

## Discussion

Our in depth analysis of *hb* DAE enhancer revealed that, strikingly, while mutations in Bcd and Zld binding sites often had at most modest effects on its activity, the mutations of the sequences between binding sites of these two factors and Gt strongly decreased its activity. This later finding suggested that sequences between the known transcription factors Zld, Bcd, and Gt are important for the activity of the DAE. This is quite surprising since this enhancer is very similar to *hb* promoter proximal enhancer (Perry et al., 2011), which is well known to be primarily activated by Bcd (Driever et al., 1989; Struhl et al., 1989), and Zld and Bcd were expected to be all that is required for *hb* DAE mediated activation.

**Fig. S4.**
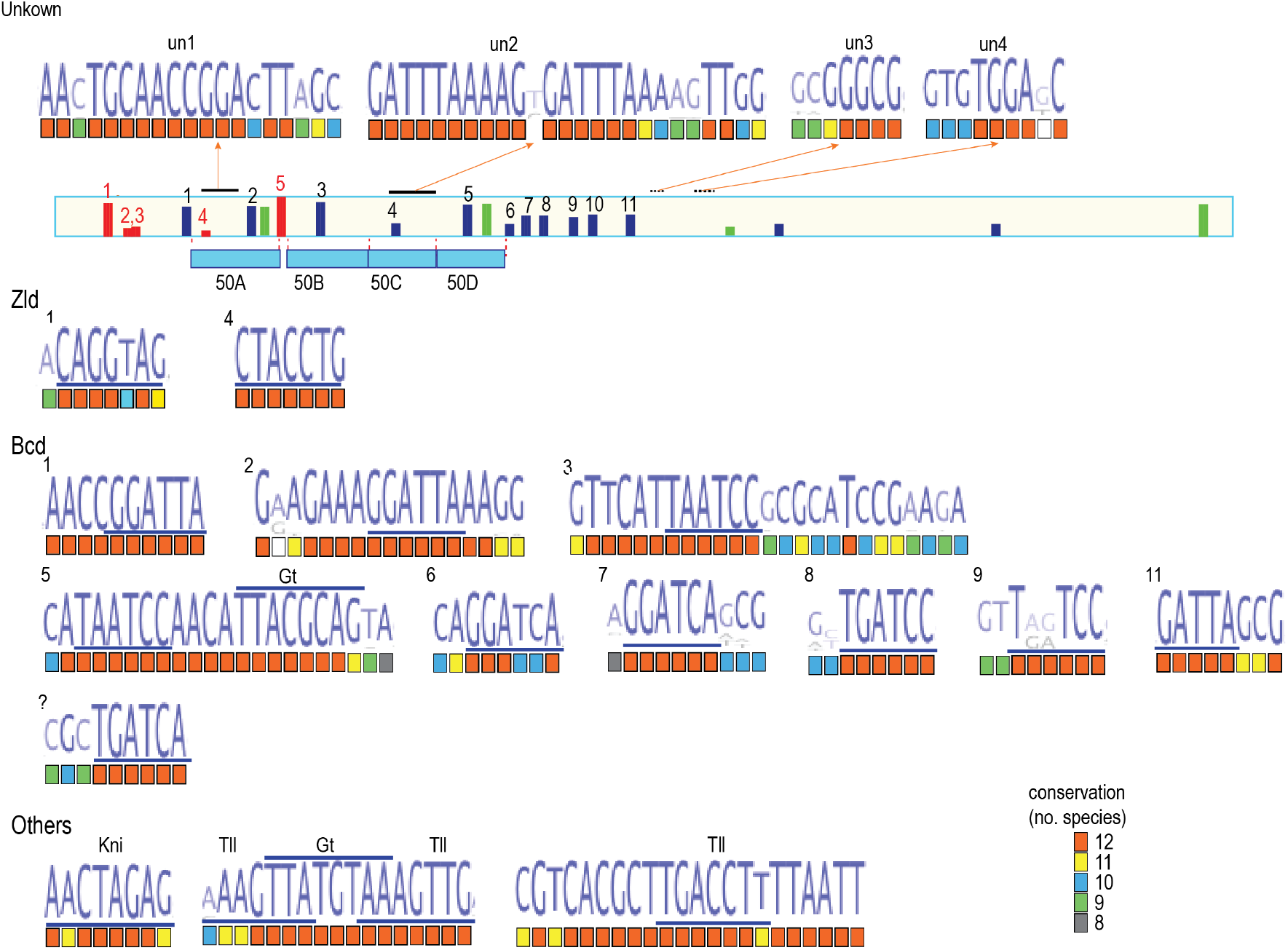
Conserved sequences in the *hb* DAE based on sequence alignment between different *Drosophila* species. *hb* DAE sequences from twelve *Drosophila* species were aligned using the multiple sequence alignment program MUSCLE from EMBL-EBI. The CLUSTAL output was loaded onto jalview and adjustments of alignment were made to achieve best align for the Zld and Bcd sites between the species. Among the motifs (underlined) for Zld, Bcd, Gt, and Til identified by patser from *D. melanogaster*, all except Zld sites 3 and 4, and Bcd sites 4 and 9 were conserved. The conserved sequences flanking the factor motifs are shown along with the motif sequences. Also shown are the conserved blocks (un1-4) that did not match a known motif in the fly factor database. The color in the square under each base denotes the number of species in which the base is conserved

**Fig. S5.**
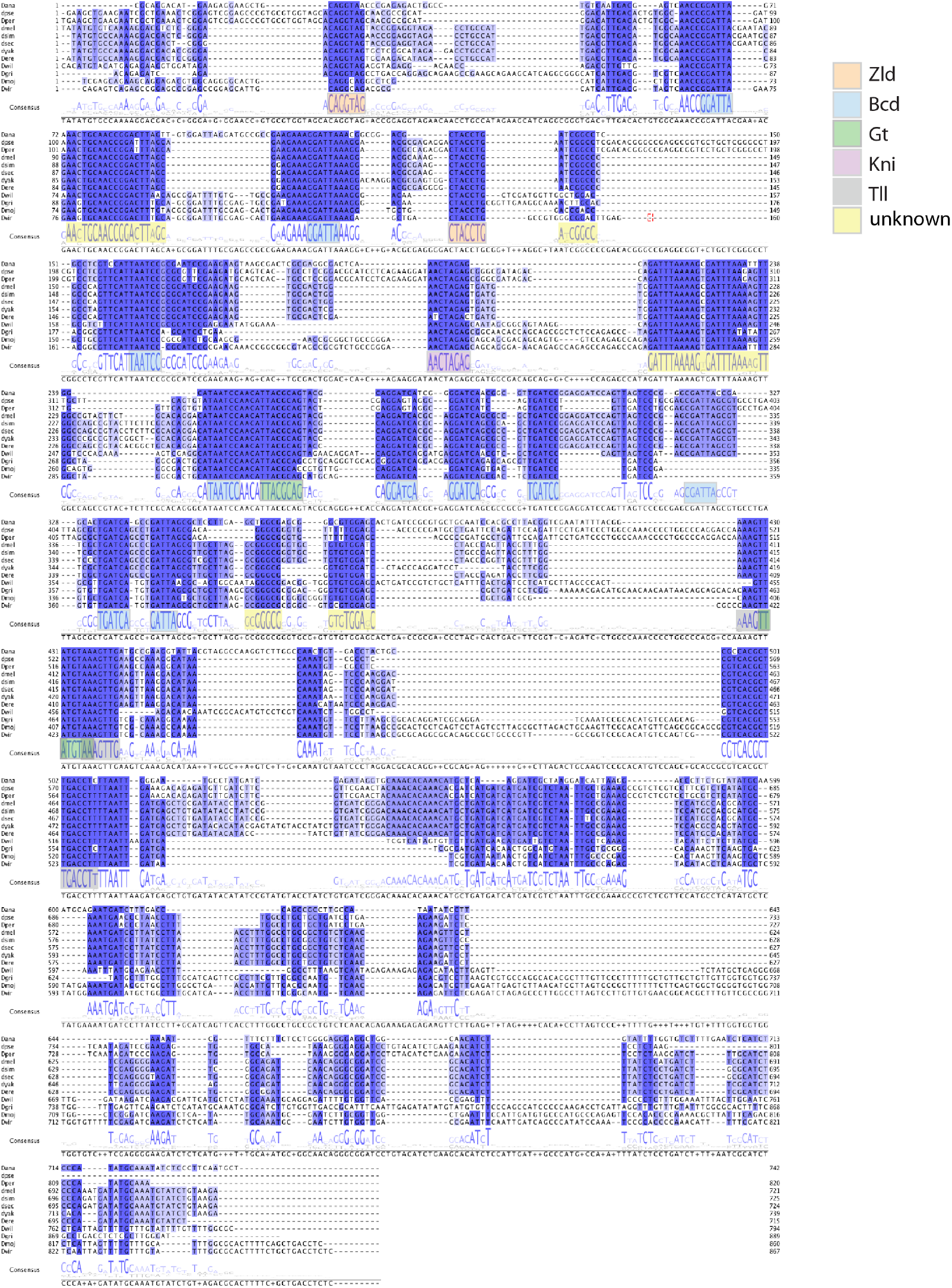
Alignment of the *hb* DAE enhancer sequences from twelve Drosophila species. *hb* DAE sequences from twelve *Drosophila* species were aligned as described in Fig.S4. The species are indicated on the left. The conserved sequence blocks are marked by different colors.

One possible explanation for the loss of DAE activity upon mutations of the sequences between Zld, Bcd, and Gt sites is that those sequences contain the binding sites of a certain unknown factor(s). We consider this unlikely for several reasons. First of all, such factor needs to be expressed broadly to at least cover the anterior half of the embryo where the DAE is active. Within the transcription network of the early fly embryo, only *hb* itself meets this expression pattern criteria, and can function as an activator (Small et al., 1991). However, the DAE does not contain *hb* binding sites and is not bound significantly by *hb* based on ChIP data (e.g. (Li et al., 2008)). In addition, the Gaf factor is expressed ubiquitously and its binding motif is enriched among enhancer sequences (Yáñez-Cuna et al., 2014). However, *hb* DAE does not contain the GAGA motif for the Gaf factor either. We also tried to find potential transcription factor binding sites by finding conserved sequences outside Zld, Bcd, and Gt sites, and matching them known motifs in the fly factor database, but we failed to find any except for matches to known repressors, Kni and Tll, which may modulate the activity of this enhancer through repression. The conserved sequences identified by the alignment analysis are also very diverse, and none of them was present at more than one location in the DAE to be able to explain the broad requirement for the sequences between the known transcription factors revealed in the mutational analyses. Take all this consideration, we favor the possibility that the sequences between the transcription factor binding sites do not provide binding site information for transcription factors, but instead may play some other functional roles in determining the DAE activity.

Even though, our finding that the sequences between transcription factor binding sites are broadly required for enhancer activity is quite surprising, some previous studies have indicated that non-transcription factor binding sequences can be important and can affect enhancer activity in various ways. One way the enhancer sequences can affect enhancer activity is by influencing nucleosome occupany and positioning. In this regard, it is known that yeast promoters are often associated with the nucleosome-destabilizing poly(dA-dT) track, which is important for transcriptional activation in yeast (Iyer and Struhl, 1995; Raveh-Sadka et al., 2012). In fly and mammals, the enhancers are generally associated with high nucleosome occupancy (Tillo and Hughes, 2009), which may be related to the high GC content of enhancer sequences (Fenouil et al., 2012; Li et al., 2008; Thurman et al., 2012). The high nucleosome occupancy is likely to have functional consequences for enhancer activity. Interestingly, it was found that for the hematopoietic pioneer factor Pu.1 bound enhancers, the DNA sequence and shape features that determine the binding of this factor and the nucleosome occupancy are highly correlated (Barozzi et al., 2014), which also suggests functional role of sequences between transcription factors.

Some other studies have shown that transcription factor binding affinity and specificity can be affected by DNA shape of the sequences around the transcription factor binding sites (Gordân et al., 2013) (Slattery et al., 2014). This may represent another important way sequences outside transcription factor binding motifs can affect transcription factor binding and enhancer activity. Moreover, it has been shown recently that enhancers in *Drosophila* are enriched with dinucleotide repeats, such as CACA repeats, which are important for enhancer activity (Yáñez-Cuna et al., 2014), although how the these repetitive sequence affect enhancer activity is unclear.

For the *hb* DAE, it is conceivable that the conserved sequences flanking the Bcd sites may influence Bcd binding by affecting the local DNA shape. It is also possible that the sequences between the transcription factor binding sites may be important for nucleosome positioning which may be important for the DAE activity. In support of this, we found that quite interestingly, this enhancer has a pair of well positioned nucleosomes in the critical segment of the enhancer (Fig. S6). It is conceivable that the sequences between the transcription factor binding sites may be important for determining the nucleosome positioning, which may be important for the activity of this enhancer. In future studies it will be very interesting to find out how this nucleosome positioning is encoded in the enhancer sequence, and how it will affect the enhancer activity, and whether the sequences between the transcription factors also affects the enhancer activity through other mechanisms.

**Fig. S6.**
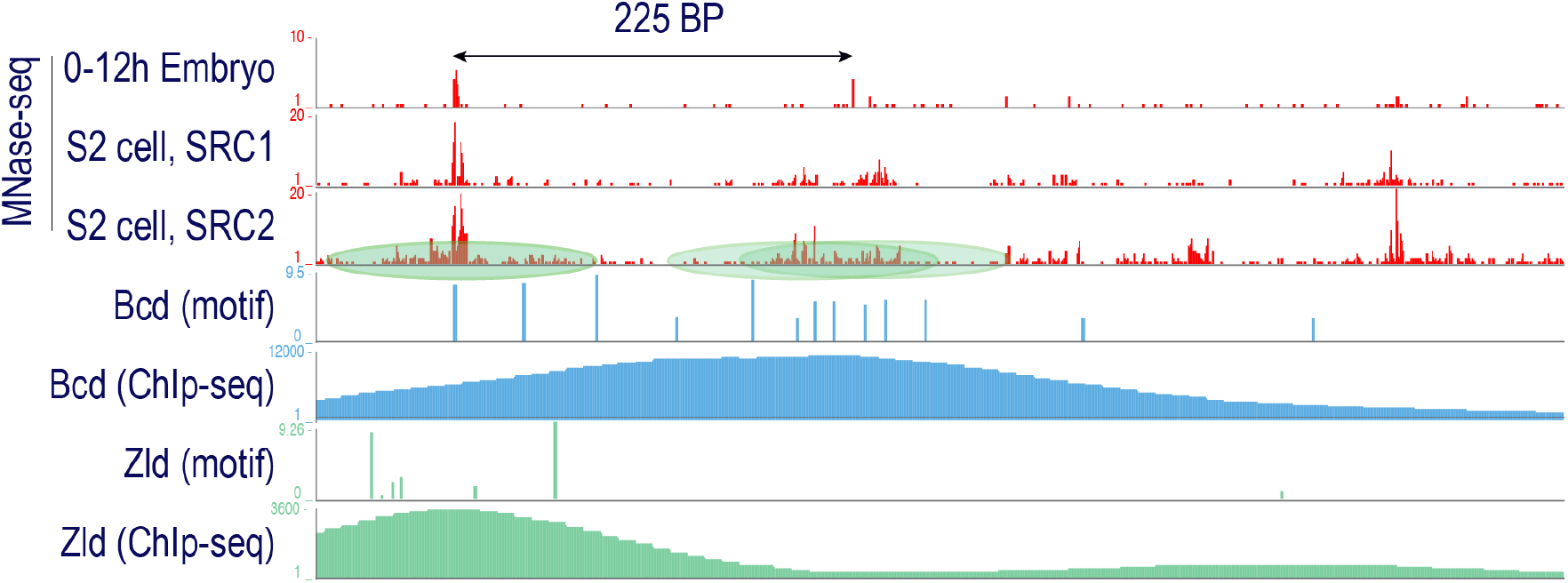
Nucleosome positioning at *hb* DAE. Distributions of aligned MNase-seq read mid-points based on three sets of MNase-seq data obtained from 0-12 hr old embryos (Chereji et al., 2016) or S2 cells (SRC1: (Gilchrist et al., 2010); SRC2: (Chereji et al., 2016)) are shown. The two positioned nucleosomes inferred from the MNase-seq read mid-point distributation are schematically shown. The Bcd (Bradley 2010) and Zld (Harrison, 2011) ChIP-seq profiles are shown based on data from previous report.

In summay, starting with an enhancer which was thought to have a simple transactivating input in the form of binding sites for Zld and Bcd, we found that the sequences between Zld and Bcd sites harbor important information required for *hb* DAE activity. While we cannot exclude the possibility that an uncharacterized transcription factor may contribute, we have enough evidence to suggest that the sequences do not function through transcription factor binding, and instead may affect *hb* DAE activity by other mechanisms. Characterizing the function of these non-factor binding sequences will be a challenge in future studies but will be important towards our full understanding of enhancer structure and function.

## Materials and Methods

### Identification of Transcription Factor Binding Sites in *hb* DAE Enhancer

The binding sites for each factor within the *hb* DAE enhancer sequence were predicted with patser (version 3e) (Hertz and Stormo, 1999) with ln(p-value) cutoffs of -6 for Bcd, and -6.5 for Gt, and Zld. The Position Weight Matrices used for Bcd and Gt (Li et al., 2008) and Zld (Enuameh et al., 2013) have been described previously.

### Creation of Mutant Enhancers

To generate random sequences used to replace the sequences between transcription factor binding sites, each random base pair was pick from the four nucleotides ACGT after shuffling them using the perl fisher yates_shuffle module. The random sequences were first scanned with patser program to make sure no new Bcd, Zld, and Gt sites were generated by chance. The mutant enhancers containing random sequence replacement between core or full transcription factor binding sites throughout the enhancer sequence, *hb* DAE(Rd) and *hb* DAE(Rd-f), and the mutant enhancer sequence in which the sequences between transcription factor binding sites were altered by swapping the top and bottom bases for each base pair were synthesized as gBlocks gene fragments (Integrated DNA Technologies). The synthesized DNA fragments also included the 50 bp sequences flanking the *eve* stripe 1 enhancer, which are necessary for swapping the mutant enhancer sequences in place of the galK sequence in *eve* bac vector by recombineering (see below).

To generate enhancer mutants in which only sequences in a certain region (200bp or 50 bp) of the enhancer were replaced with random sequences, an overlap extension PCR method (Horton et al., 1989; Yon and Fried, 1989) was used. The scheme to generate the sequence is shown in Fig.S7, along with the primers used. Briefly, to create the constructs containing random sequences in 200 bp regions (hb-DAE(Rd-200A-C)) (Fig. S7B,E), the regions containing the random sequences were amplified from the hb-DAE(Rd) mutant enhancer construct, while the wild type segments were amplified from the wild type *hb* DAE reporter construct. The primers used to amplify the mutant and the wild type segments contained overlapping sequences, allowing the overlap extension PCR to sew pairs of (or three) mutant and wild type sequences together to obtain the final mutant enhancer.

To create constructs in which only a 50 bp region was altered (Fig.S7D,E), the upstream and downstream segments of the enhancer flanking the 50 region were amplified by PCR using the wild type *hb* DAE reporter construct as template, and using the upstream and downstream primers, each of which was paired with a primer flanking the 50 bp region. The two primers flanking the 50bp regions also contained additional sequences that correspond to overlapping portions of the 50 bp mutant sequence, which allow the creation of the 50 bp mutant region after the two segments were joined by overlap extension/sewing PCR with the primers that flank the whole enhancer to create the partially altered enhancer.

The Bcd binding site mutants were also created by overlap extension PCR (Fig.S8. In this case, however, the primers overlapping the Bcd sites to be mutated were used. The mutations were introduced to the primers, which were used together with primers matching 5’ and 3’ ends of enhancer sequence to amplify segments of the enhancer sequence. The resulting fragments were then stitched together by PCR using the primers flanking the enhancer, producing the full-length mutated enhancer.

**Fig. S7.**
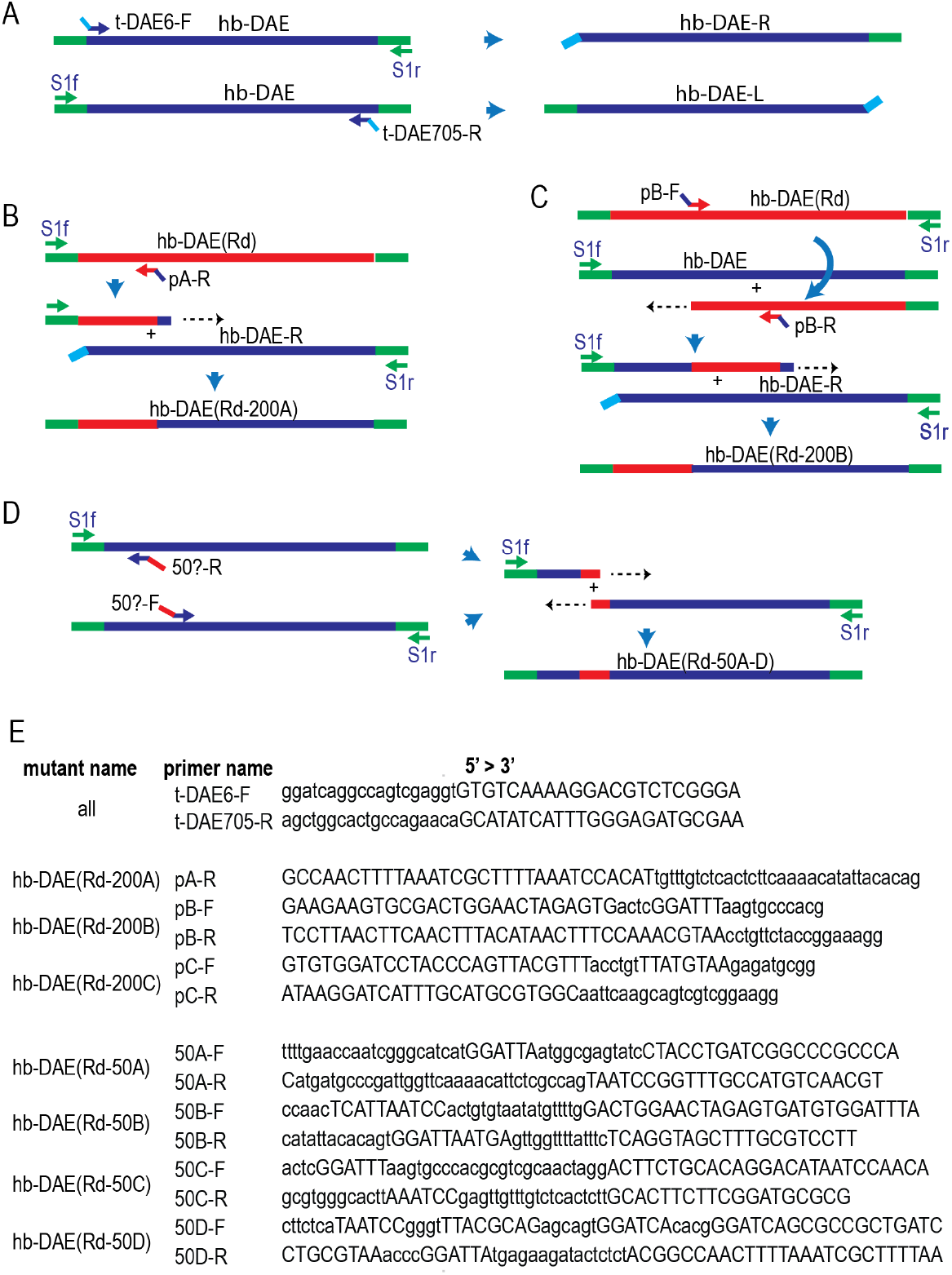
Creation of *hb* DAE mutant enhancers that carry partial sequence changes. The *hb* DAE sequence is flanked by *eve* sequences in the *hb* DAE reporter construct. It was PCR amplified to produce products that carry randomly chosen sequences either at the 5’ or 3’ end, which were added to prevent self amplification in the subsequent fusion PCR reactions. B. Scheme used to create hb-DAE(Rd-200A). C. Scheme used to create hb-DAE(Rd-200B). The hb-DAE(Rd-200C) was made in similar way. D. Scheme used to create mutant enhancer that carry sequence changes in 50 bp regions. E. List of oligos used.

**Fig. S8.**
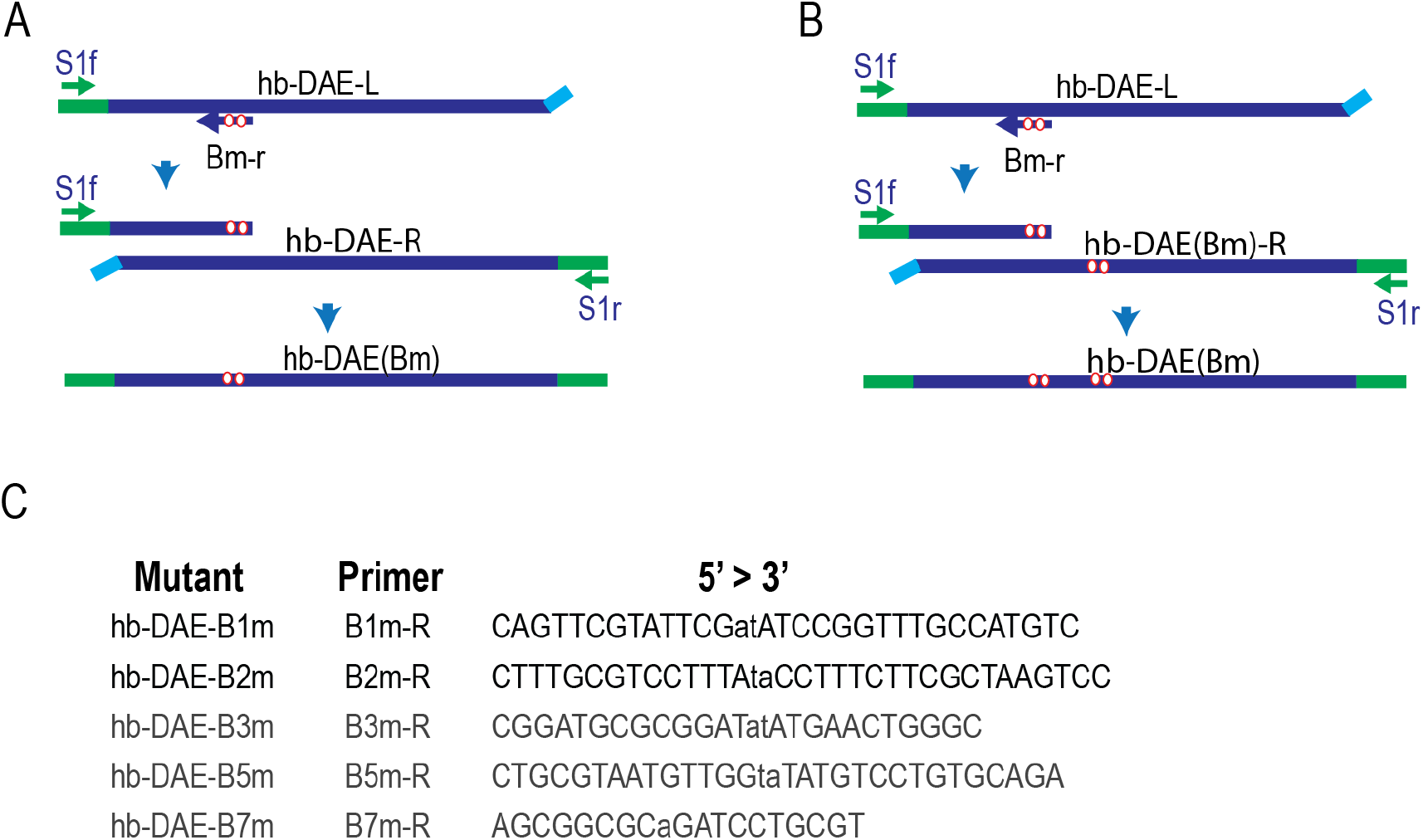
Creation of *hb* DAE Bcd binding site mutant enhancers. A. Scheme for creating single point mutations. Hb-DAE-L and hb-DAE-R were prepared as shown in Fig. S7. B. Scheme for creating double Bcd site mutants, starting from single site mutant construct. Hb-DAE-L, and hb-DAE(Bm) -R were produced through PCR amplification as described in Fig.S3 except the hb-DAE(Bm) -R was made using a single Bcd site mutant enhancer as template. C. The oligos used to introduce the each single Bcd mutation. The nucleotides changed from wild type are in lowercase in the sequences.

### Genomic sequences enriched with Bcd sites

The fly genome was scanned for presence of three or more Bcd binding sites in a 75bp window. The resulting significant sequence intervals were then merged if they were within 100 bp to generate a list of Bcd site cluster sequences. These sequences were ranked based on total weight of the Bcd sites. We selected two top ranked sequences and inserted the Zld binding site containing sequences from *hb* into these sequences in such a way so that the arrangement of the Zld sites relative to the Bcd sites are similar to those in the *hb* DAE. The sequences can be found in the supplement.

### Recombineering and Transgenic Flies

The *eve* bac reporter vector and the enhancer reporter constructs were created through recombineering (Lee et al., 2001; Liu et al., 2003; Warming et al., 2005) (Venken et al., 2009). The starting *eve* bac plasmid P[acman]-CH322-103k22-GFP, which contains the eGFP-Kan^r^ tag sequence at the C-terminus of the *eve* CDS, was previously described (Venken et al., 2009). The starting *eve* bac plasmid P[acman]-CH322-103k22-GFP, We removed the CDS of the *eve* gene in this construct to generate peve-GFP by standard recombineering procedure, by first replacing the *eve* CDS with galK sequence amplified from a galK plasmid pBALU1 using primers flanked by *eve* sequences:

eve-galKf: 5’ TTAATATCCTCTGAATAAGCCAACTTTGAATCACAAGACGCATACCAAACCCT GTT GACAATTAATCATCGGCA 3’

eve-galKr: 5’ GCGCCCTGAAAATAAAGATTCTCGCTTGCAGTAGTTGGAATATCATAATCTC AGCACTGTCCTGCTCCTT 3’.

Then the galk sequence was replaced with:

5’ ATATCCTCTGAATAAGCCAACTTTGAATCACAAGACGCATACCAAACATGGATTATGAT ATTCCAACTACTGCAAGCGAGAATCTTTATTTTCAGGGCGC 3’

We further replaced the *eve* stripe 2 and then *eve* stripe 1 enhancers in this plasmid with bacteria amp^r^ gene and the bacteria galK gene respectively to generate the *eve* bac reporter vector, peve-GFP/S2xAmp^r^-S1xgalK. To replace of *eve* stripe 2 sequence with amp^r^ sequence, the sequence was first replaced with galK sequence using PCR product amplified from the galK containing plasmid with:

eve-S2-galKf: 5’ AGGGATTCGCCTTCCCATATTCCGGGTATTGCCGGCCCGGGAAAATGCGACCTGTTGACAATTAATCATCGGCA 3’

eve-S2-galKr: 5’ TTGCGCAAGTTAGCTTGGAGGTTTGGCCAAAAAAATCGTGGGGTCCACCCTCAGCACTGTCCTGCTCCTT 3’

Then, the galK sequence was replaced with amp^r^ sequence amplified by PCR from an amp^r^ containing vector plasmid using:

Eve-S2-Ampf: 5’ AGGGATTCGCCTTCCCATATTCCGGGTATTGCCGGCCCGGGAAAATGCGAGGGGTCTGACGCTCAGTGGAACGAA 3’

Eve-S2-Ampr: 5’ TCGGTTTCGTTGGGCAAAACATTTATTATGATGATATAATCATTCCC AGCGTGGCACTTTTCGGGGAAATGTGC 3’

The *eve* S1 sequence was replaced with galK sequence amplified by PCR with:

Eve-S1-galKf: 5’ TACGGATCGGAATTGGGATCGAAATCGGGATATGCACAACAC GGTAATGACCTGTTGACAATTAATCATCGGCA 3’

Eve-S1-galKr: 5’ AACACATCGCCGGACAAAGGGAAATTTGCTGCCGACTTGTTG ACTTTGCGTCAGCACTGTCCTGCTCCTT 3’

All other enhancer reporter constructs were created by swapping out the galK sequence in bac vector generated above through recombineering and replaced with the wild type and mutant *hb* DAE enhancer DNA either custom made (Integrated DNA technologies) or by PCR as described above. These DNA sequences all were flanked by sequences around the *eve* stripe 1 sequence that was replaced by the galK sequence as underlined in the two oligos listed above.

The transgenes were generated by injecting the reporter construct plasmids into the embryos from flies that carry the VK33 (Venken et al., 2006) attp landing site on chromosome 3 using the services of Rainbow Transgenic flies Inc and Bestgene Inc. The transgenes were identified by crossing the G0 flies from injected embryos to a yw; TM3 sb balancer line and screening for red eye phenotype. Homozygous lines for the transgenes were established by selfing the balanced red eye flies.

### Florescence in situ Hybridization and Embryo Imaging

FISH was carried out with embryos collected from the transgene flies and fixed with formaldehyde as previously described (Kosman et al., 2004) with some modifications.

To prepare the RNA probe for the reporter gene, the eGFP reporter gene and the kan^R^ gene sequences were amplified by PCR separately from the *eve* reporter construct using the following oligos:

eGFP-F: 5’ GGATTATGATATTCCaACTACTGCAAGCGAG 3’

T7p-eGFP-R: 5’

CAGGTCTGAGTAATACGACTCACTATAGGGTACAGCTCGTCCATGCC GAG 3’

Kan-F: 5’ GTTACTGGCCGAAGCCGCTTG 3’

T7p-Kan-R: 5’

CAGGTCTGAGTAATACGACTCACTATAGGGAAGAACTCGTCAAGAAGGC GATAGAAG 3’

The reverse primers used contained T7 promoter sequence at their 5’ end, allowing the PCR products to be used directly for RNA probe synthesis. The probes were made using Thermo Transcript Aid T7 High Yield Transcription Kit (ThermoFisher Scientific, cat. no. K0441) with the Digoxigenin(DIG) RNA labeling mix from Roche (cat. no. 1277073) and were not hydrolyzed before being used.

The embryos were collected for 2 hours and aged for 1 hr and 20 min and were fixed as described in (Kosman et al. 2004). The preparation before hybridization, and the hybridization step were carried out as previously described, except that the protease K treatment and the second post-fix step that followed were skipped. For florescent detection, the embryos were incubated with pre-absorbed sheep anti-DIG-POD (Roche, cat. no. 11207733910). Following incubation and washing, the embryos were mixed and incubated with TSA coumarin in amplification buffer and incubated by rocking. The embryos were washed, and near the end of wash were incubated with Sytox Green nucleic acid stain (ThermoFisher Scienctific, cat. no. S7020) to label the nuclei. The embryos were washed again once, and mounted in 70% glycerol, 2.5% DABCO.

The embryo imaging was performed on a Zeiss LSM800 microscope with a 10x objective. For all experiments, Z stack images were taken from top to the mid-section of each embryo with the image size set at 2014×849, or a resolution of about 0.27 μm/pixel. The other settings differed between experiments, which was not anticipated to affect the results since reporter activities detected for the mutant enhancers were always compared to the signals detected for wild type enhancer in the same experiment. For earlier experiments, 15 z stacks were taken for each embryo with thickness of about 5.6 μm, and the fluorescence signals for coumarin with an excitation wavelength of 405 and Sytox Green with excitation wavelength at 488 were obtained sequentially with the “smart” setting. For later experiments, 30 z stacks of images per embryo were taken, and the two flurorence signals were obtained simultaneously with the “fast” setting. In all cases, the laser power and digital gain for each dye were adjusted in each experiment to make sure the maximum signal obtained was below saturation.

The images were analyzed with the Zeiss image analysis software zen 2. A 2-D image was generated for each embryo from the z stack images using the orthogonal maximum projection function. Then the fluorescent signal profiles for the RNA (coumarin) and nuclei (sytox green) were generated for a rectangle area drawn to the length of the embryo along the A/P axis, and with a width of 200 pixels, about a third of the embryo width. The signals along the A/P axis were then divided into 100 bins corresponding to the percent egg length. The coumarin signals were then divided by the corresponding Sytox signals as a simple way of normalization to correct for the nuclear density variation between different part of the embryo and between embryos. Furthermore, since the reporter signal between egg length 50% - 60% is the lowest in the coumarin signal profile, and not affected by the presence *hb* DAE, and the average of signal in this region was taken as background and subtracted from the signal throughout the whole embryo.

## Reference

Arnosti, D.N., Barolo, S., Levine, M., and Small, S. (1995). The *eve* stripe 2 enhancer employs multiple modes of transcriptional synergy. Development 122, 205–214.

Barozzi, I., Simonatto, M., Bonifacio, S., Yang, L., Rohs, R., Ghisletti, S., and Natoli, G. (2014). Coregulation of Transcription Factor Binding and Nucleosome Occupancy through DNA Features of Mammalian Enhancers. Molecular Cell 54, 844–857.

Chereji, R.V., Kan, T.-W., Grudniewska, M.K., Romashchenko, A.V., Berezikov, E., Zhimulev, I.F., Guryev, V., Morozov, A.V., and Moshkin, Y.M. (2016). Genome-wide profiling of nucleosome sensitivity and chromatin accessibility in Drosophila melanogaster. Nucleic Acids Research 44, 1036–1051.

Crocker, J., Tsai, A., and Stern, D.L. (2017). A Fully Synthetic Transcriptional Platform for a Multicellular Eukaryote. Cell Rep 18, 287–296.

Driever, W., and Nüsslein-Volhard, C. (1988). A gradient of bicoid protein in Drosophila embryos. Cell 54, 83–93.

Driever, W., Thoma, G., and Nüsslein-Volhard, C. (1989). Determination of spatial domains of zygotic gene expression in the Drosophila embryo by the affinity of binding sites for the bicoid morphogen. Nature 340, 363–367.

Driever, W., and Nüsslein-Volhard, C. (1989). The bicoid protein is a positive regulator of hunchback transcription in early drosophila embryo. Nature 337, 138143.

Enuameh, M.S., Asriyan, Y., Richards, A., Christensen, R.G., Hall, V.L., Kazemian, M., Zhu, C., Pham, H., Cheng, Q., Blatti, C., et al. (2013). Global analysis of Drosophila Cys2-His2 zinc finger proteins reveals a multitude of novel recognition motifs and binding determinants. Genome Research 23, 928–940.

Farley, E.K., Olson, K.M., zhang, W., Brandt, A.J., Rokhsar, D.S., and Levine, M.S. (2015). Suboptimization of developmental enhancers. Science 350, 325–328.

Fenouil, R., Cauchy, P., Koch, F., Descostes, N., Cabeza, J.Z., Innocenti, C., Ferrier, P., Spicuglia, S., Gut, M., Gut, I., et al. (2012). CpG islands and GC content dictate nucleosome depletion in a transcription-independent manner at mammalian promoters. Genome Research 22, 2399–2408.

Fujioka, M., Mistry, H., Schedl, P., and Jaynes, J.B. (2016). Determinants of Chromosome Architecture: Insulator Pairing in cis and in trans. PLoS Genetics 12, e1005889.

Gilchrist, D.A., Santos, Dos G., Fargo, D.C., Xie, B., Gao, Y., Li, L., and Adelman, K. (2010). Pausing of RNA polymerase II disrupts DNA-specified nucleosome organization to enable precise gene regulation. Cell 143, 540–551.

Gordân, R., Shen, N., Dror, I., Zhou, T., Horton, J., Rohs, R., and Bulyk, M.L. (2013). Genomic regions flanking E-box binding sites influence DNA binding specificity of bHLH transcription factors through DNA shape. Cell Rep 3, 1093–1104.

Goto, T., Macdonald, P., and Maniatis, T. (1989). Early and late periodic patterns of even skipped expression are controlled by distinct regulatory elements that respond to different spatial cues. Cell 57, 413–422.

Hannon, C.E., Blythe, S.A., and Wieschaus, E.F. (2017). Concentration dependent chromatin states induced by the bicoid morphogen gradient. eLife 6.

Harrison, M.M., and Eisen, M.B. (2015). Transcriptional Activation of the Zygotic Genome in Drosophila (Elsevier Inc.).

Harrison, M.M., Li, X.-Y., Kaplan, T., Botchan, M.R., and Eisen, M.B. (2011). Zelda binding in the early Drosophila melanogaster embryo marks regions subsequently activated at the maternal-to-zygotic transition. PLoS Genetics 7, e1002266.

Hertz, G.Z., and Stormo, G.D. (1999). Identifying DNA and protein patterns with statistically significant alignments of multiple sequences. Bioinformatics 15, 563577.

Horton, R.M., Hunt, H.D., Ho, S.N., Pullen, J.K., and Pease, L.R. (1989). Engineering hybrid genes without the use of restriction enzymes: gene splicing by overlap extension. Gene 77, 61–68.

Iyer, V., and Struhl, K. (1995). Poly(dA:dT), a ubiquitous promoter element that stimulates transcription via its intrinsic DNA structure. The EMBO Journal 14, 2570–2579.

Iyer, V.R., Horak, C.E., Scafe, C.S., Botstein, D., Snyder, M., and Brown, P.O. (2001). Genomic binding sites of the yeast cell-cycle transcription factors SBF and MBF. Nature 409, 533–538.

Jolma, A., Yin, Y., Nitta, K.R., Dave, K., Popov, A., Taipale, M., Enge, M., Kivioja, T., Morgunova, E., and Taipale, J. (2015). DNA-dependent formation of transcription factor pairs alters their binding specificity. Nature 527, 384–388.

Kosman, D., Mizutani, C.M., Lemons, D., Cox, W.G., McGinnis, W., and Bier, E. (2004). Multiplex detection of RNA expression in Drosophila embryos. Science 305, 846.

Lee, E.C., Yu, D., Martinez de Velasco, J., Tessarollo, L., Swing, D.A., Court, D.L., Jenkins, N.A., and Copeland, N.G. (2001). A highly efficient Escherichia coli-based chromosome engineering system adapted for recombinogenic targeting and subcloning of BAC DNA. Genomics 73, 56–65.

Lelli, K.M., Slattery, M., and Mann, R.S. (2012). Disentangling the many layers of eukaryotic transcriptional regulation. Annu. Rev. Genet. 46, 43–68.

Levine, M. (2010). Transcriptional enhancers in animal development and evolution. Curr. Biol. 20, R754–R763.

Levo, M., zalckvar, E., Sharon, E., Dantas Machado, A.C., Kalma, Y., Lotam-Pompan, M., Weinberger, A., Yakhini, Z., Rohs, R., and Segal, E. (2015). Unraveling determinants of transcription factor binding outside the core binding site. Genome Research 25, 1018–1029.

Li, X.-Y., MacArthur, S., Bourgon, R., Nix, D., Pollard, D.A., Iyer, V.N., Hechmer, A., Simirenko, L., Stapleton, M., Luengo Hendriks, C.L., et al. (2008). Transcription factors bind thousands of active and inactive regions in the Drosophila blastoderm. PLoS Biology 6, e27.

Liang, H., Lin, Y.S., and Li, W.H. (2008). Fast Evolution of Core Promoters in Primate Genomes. Molecular Biology and Evolution 25, 1239–1244.

Liu, P., Jenkins, N.A., and Copeland, N.G. (2003). A highly efficient recombineering-based method for generating conditional knockout mutations. Genome Research 13, 476–484.

Liu, X., Lee, C.-K., Granek, J.A., Clarke, N.D., and Lieb, J.D. (2006). Whole-genome comparison of Leu3 binding in vitro and in vivo reveals the importance of nucleosome occupancy in target site selection. Genome Research 16, 1517–1528.

MacArthur, S., Li, X.-Y., Li, J., Brown, J.B., Chu, H.C., Zeng, L., Grondona, B.P., Hechmer, A., Simirenko, L., Keränen, S.V.E., et al. (2009). Developmental roles of 21 Drosophila transcription factors are determined by quantitative differences in binding to an overlapping set of thousands of genomic regions. Genome Biology 10, R80.

Mann, R.S., Lelli, K.M., and Joshi, R. (2009). Hox specificity unique roles for cofactors and collaborators. Curr. Top. Dev. Biol. 88, 63–101.

Miller, J.A., and Widom, J. (2003). Collaborative competition mechanism for gene activation in vivo. Mol. Cell. Biol. 23, 1623–1632.

Mirny, L.A. (2010). Nucleosome-mediated cooperativity between transcription factors. Pnas 107, 22534–22539.

Panne, D., Maniatis, T., and Harrison, S.C. (2007). An Atomic Model of the Interferon-β Enhanceosome. Cell 129, 1111–1123.

Perry, M.W., Boettiger, A.N., and Levine, M. (2011). Multiple enhancers ensure precision of gap gene-expression patterns in the Drosophila embryo. Pnas 108, 13570–13575.

Raveh-Sadka, T., Levo, M., Shabi, U., Shany, B., Keren, L., Lotan-Pompan, M., Zeevi, D., Sharon, E., Weinberger, A., and Segal, E. (2012). Manipulating nucleosome disfavoring sequences allows fine-tune regulation of gene expression in yeast. Nature Publishing Group 44, 743–750.

Slattery, M., Zhou, T., Yang, L., Machado, A.C.D., Gordân, R., and Rohs, R. (2014). Absence of a simple code: howtranscription factors read the genome. Trends in Biochemical Sciences 39, 381–399.

Small, S., Blair, A., and Levine, M. (1992). Regulation of even-skipped stripe 2 in the Drosophila embryo. The EMBO Journal 11, 4047–4057.

Small, S., Kraut, R., Hoey, T., Warrior, R., and Levine, M. (1991). Transcriptional regulation of a pair-rule stripe in Drosophila. Genes & Development 5, 827–839.

Spitz, F., and Furlong, E.E.M. (2012). Transcription factors: from enhancer binding to developmental control. Nature Reviews Genetics 13, 613–626.

Stanojevic, D., Small, S., and Levine, M. (1991). Regulation of a segmentation stripe by overlapping activators and repressors in the Drosophila embryo. Science 254, 1385–1387.

Struhl, G., Struhl, K., and Macdonald, P.M. (1989). The gradient morphogen bicoid is a concentration-dependent transcriptional activator. Cell 57, 1259–1273.

Tautz, D. (1988). Regulation of the Drosophila segmentation gene hunchback by two maternal morphogenetic centres. Nature 332, 281–284.

Thurman, R.E., Rynes, E., Humbert, R., Vierstra, J., Maurano, M.T., Haugen, E., Sheffield, N.C., Stergachis, A.B., Wang, H., Vernot, B., et al. (2012). The accessible chromatin landscape of the human genome. Nature 489, 75–82.

Tillo, D., and Hughes, T.R. (2009). G+C content dominates intrinsic nucleosome occupancy. BMC Bioinformatics 10, 442.

Venken, K.J.T., Carlson, J.W., Schulze, K.L., Pan, H., He, Y., Spokony, R., Wan, K.H., Koriabine, M., de Jong, P.J., White, K.P., et al. (2009). Versatile P[acman] BAC libraries for transgenesis studies in Drosophila melanogaster. Nature Methods 6, 431–434.

Venken, K.J.T., He, Y., Hoskins, R.A., and Bellen, H.J. (2006). P[acman]: a BAC transgenic platform for targeted insertion of large DNA fragments in D. melanogaster. Science 314, 1747–1751.

Vincent, B.J., Estrada, J., and DePace, A.H. (2016). The appeasement of Doug: a synthetic approach to enhancer biology. Integr Biol (Camb) 8, 475–484.

Warming, S., Costantino, N., Court, D.L., Jenkins, N.A., and Copeland, N.G. (2005). Simple and highly efficient BAC recombineering using galK selection. Nucleic Acids Research 33, e36.

Wunderlich, Z., and Mirny, L.A. (2009). Different gene regulation strategies revealed by analysis of binding motifs. Trends in Genetics : TIG 25, 434–440.

Xu, Z., Chen, H., Ling, J., Yu, D., Struffi, P., and Small, S. (2014). Impacts of the ubiquitous factor Zelda on Bicoid-dependent DNA binding and transcription in Drosophila. Genes & Development 28, 608–621.

Yang, A., Zhu, Z., Kapranov, P., McKeon, F., Church, G.M., Gingeras, T.R., and Struhl, K. (2006). Relationships between p63 binding, DNA sequence, transcription activity, and biological function in human cells. Molecular Cell 24, 593–602.

Yáñez-Cuna, J.O., Arnold, C.D., Stampfel, G., Boryń, L.M., Gerlach, D., Rath, M., and Stark, A. (2014). Dissection of thousands of cell type-specific enhancers identifies dinucleotide repeat motifs as general enhancer features. Genome Research 24, 1147–1156.

Yon, J., and Fried, M. (1989). Precise gene fusion by PCR. Nucleic Acids Research 17, 4895.

Zaret, K.S., and Carroll, J.S. (2011). Pioneer transcription factors: establishing competence for gene expression. Genes & Development 25, 2227–2241.

